# The house spider genome reveals an ancient whole-genome duplication during arachnid evolution

**DOI:** 10.1101/106385

**Authors:** Evelyn E. Schwager, Prashant P. Sharma, Thomas Clarke, Daniel J. Leite, Torsten Wierschin, Matthias Pechmann, Yasuko Akiyama-Oda, Lauren Esposito, Jesper Bechsgaard, Trine Bilde, Alexandra D. Buffry, Hsu Chao, Huyen Dinh, HarshaVardhan Doddapaneni, Shannon Dugan, Cornelius Eibner, Cassandra G. Extavour, Peter Funch, Jessica Garb, Luis B. Gonzalez, Vanessa L. Gonzalez, Sam Griffiths-Jones, Yi Han, Cheryl Hayashi, Maarten Hilbrant, Daniel S.T. Hughes, Ralf Janssen, Sandra L. Lee, Ignacio Maeso, Shwetha C. Murali, Donna M. Muzny, Rodrigo Nunes da Fonseca, Christian L. B. Paese, Jiaxin Qu, Matthew Ronshaugen, Christoph Schomburg, Anna Schönauer, Angelika Stollewerk, Montserrat Torres-Oliva, Natascha Turetzek, Bram Vanthournout, John H. Werren, Carsten Wolff, Kim C. Worley, Gregor Bucher, Richard A. Gibbs, Jonathan Coddington, Hiroki Oda, Mario Stanke, Nadia A. Ayoub, Nikola-Michael Prpic, Jean-Frangois Flot, Nico Posnien, Stephen Richards, Alistair P. McGregor

**Author notes:** Equal contribution (Evelyn E. Schwager, Prashant P. Sharma, Thomas Clarke, Daniel J. Leite, Torsten Wierschin).

## Abstract

The duplication of genes can occur through various mechanisms and is thought to make a major contribution to the evolutionary diversification of organisms. There is increasing evidence for a large-scale duplication of genes in some chelicerate lineages including two rounds of whole genome duplication (WGD) in horseshoe crabs. To investigate this further we sequenced and analyzed the genome of the common house spider *Parasteatoda tepidariorum.* We found pervasive duplication of both coding and non-coding genes in this spider, including two clusters of Hox genes. Analysis of synteny conservation across the *P. tepidariorum* genome suggests that there has been an ancient WGD in spiders. Comparison with the genomes of other chelicerates, including that of the newly sequenced bark scorpion *Centruroides sculpturatus*, suggests that this event occurred in the common ancestor of spiders and scorpions and is probably independent of the WGDs in horseshoe crabs. Furthermore, characterization of the sequence and expression of the Hox paralogs in *P. tepidariorum* suggests that many have been subject to neofunctionalization and/or subfunctionalization since their duplication, and therefore may have contributed to the diversification of spiders and other pulmonate arachnids.

## Introduction

Gene duplication plays an important role in the evolutionary diversification of organisms (Ohno, 1970; Semon and Wolfe, 2007). Unequal crossing-over commonly results in one or a few tandemly duplicated genes, but larger scale events including whole genome duplications (WGDs) can also occur. Tandem duplication has been shown to underlie the evolution of many genes in both plants and animals, for example up to 32% of genes in the centipede *Strigamia maritima* (Chipman et al., 2014; Yun et al., 2015). WGD is arguably the most sudden and massive change that a genome can experience in a single evolutionary event. The occurrence of WGDs across a wide variety of eukaryotic groups, including plants (Fawcett et al., 2009; Li et al., 2015), fungi (Ma et al., 2009; Wolfe and Shields, 1997), ciliates (Aury et al., 2006), oomycetes (Martens and Van de Peer, 2010) and animals (Amores et al., 1998; Bisbee et al., 1977; Edger and Pires, 2009; Flot et al., 2013; Jaillon et al., 2004; Nossa et al., 2014; Session et al., 2016), attest to the major impact that polyploidization events have had in reshaping the genomes of many different organisms.

Although most of the duplicated genes resulting from tandem duplication or WGD are subsequently lost, it is thought that these events provide new genetic material for some paralogous genes to undergo subfunctionalization or neofunctionalization and thus contribute to the rewiring of gene regulatory networks, morphological innovations and, ultimately, organismal diversification (Davis and Petrov, 2005; Force et al., 1999; Kellis et al., 2004; Lynch and Conery, 2000; Lynch and Force, 2000; Lynch et al., 2001; Putnam et al., 2008; Semon and Wolfe, 2007; Wolfe and Shields, 1997). Comparisons of independent paleopolyploidization events across different eukaryotes such as plants, yeast and vertebrates (Amores et al., 1998; Fawcett et al., 2009; Flot et al., 2013; Ma et al., 2009; Nossa et al., 2014; Putnam et al., 2008), have led to the development of models to try to explain genome-wide evolutionary patterns of differential gene loss and retention compared to smaller scale events (Hakes et al., 2007; Semon and Wolfe, 2007). However, the enormous differences between these disparate eukaryotic lineages in terms of genome structure, morphological and developmental organization, and ecology have impeded a critical assessment of the potential selective advantages and actual evolutionary consequences of WGDs. Thus, the extent to which WGDs may have contributed to taxonomic “explosions” and evolutionary novelties remains controversial, especially in the case of vertebrates (Donoghue and Purnell, 2005; Furlong and Holland, 2002; Hurley et al., 2007). For example, the two WGDs shared by all vertebrates have given rise to four clusters of Hox genes, which provided new genetic material that may underlie the evolutionary success and innovations among these animals (Dehal and Boore, 2005; McGinnis and Krumlauf, 1992; Putnam et al., 2008). However, only three WGD events have been demonstrated in animals other than vertebrates, one in bdelloid rotifers and possibly two in horseshoe crabs (Flot et al., 2013; Kenny et al., 2016; Nossa et al., 2014) and these events are not associated with any bursts of diversification (Ricci, 1987; Rudkin and Young, 2009). It is clear, therefore, that documenting additional examples of WGD in metazoans would significantly increase our understanding of the genomic and morphological consequence of these events.

Intriguingly, there is increasing evidence for extensive gene duplication among chelicerates other than horseshoe crabs, particularly in spiders and scorpions (Clarke et al., 2015; Clarke et al., 2014; Di et al., 2015; Fuzita et al., 2015; Fuzita et al., 2016; Janssen et al., 2010; Leite et al., 2016; Schwager et al., 2007; Sharma et al., 2015; Sharma et al., 2014b; Turetzek et al., 2016), indicating that large-scale gene duplications occurred during the evolution of these arachnids. However, although the genomes of some arachnids have been sequenced, including the tick *Ixodes scapularis* (Gulia-Nuss et al., 2016; Lawson et al., 2009), the mite *Tetranychus urticae* (Grbic et al., 2011), the Chinese scorpion *Mesobuthus martensii* (Cao et al., 2013), and two spiders - the velvet spider *Stegodyphus mimosarum* (Sanggaard et al., 2014) and the Brazilian whiteknee tarantula *Acanthoscurria geniculata* (Sanggaard et al., 2014) - a systematic analysis of genome evolution among these diverse animals has yet to be carried out (Fig. 1) (Schwager et al., 2015).

**Figure 1.**
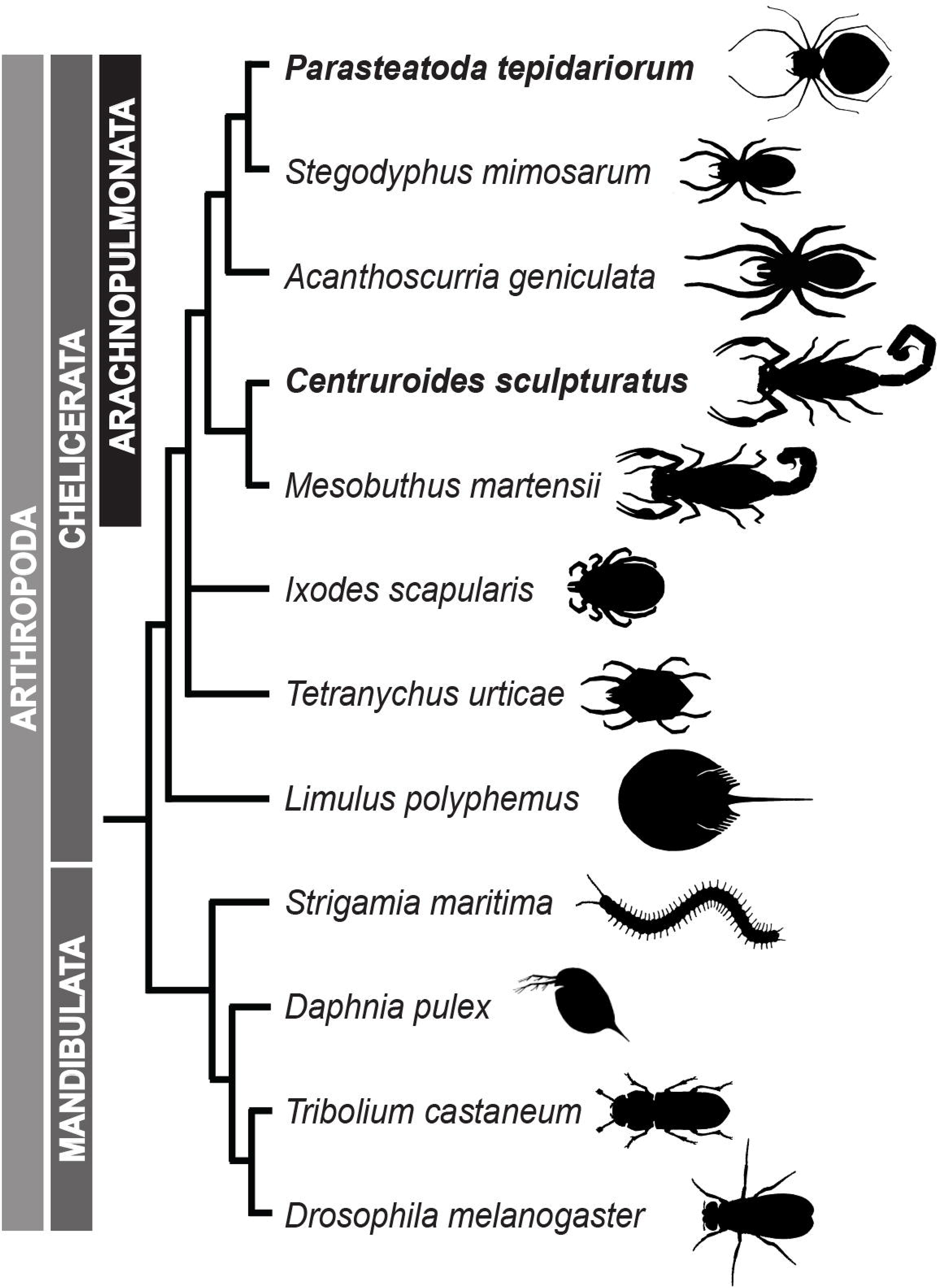
The relationships of *Parasteatoda tepidariorum* to select arthropods. Representatives of spiders (Araneae) with sequenced genomes (*P. tepidariorum, Stegodyphus mimosarum* and *Acanthoscurria geniculata*) are shown with respect to other completely sequenced chelicerates including scorpions (*Centruroides sculpturatus* and *Mesobuthus martensii)*, a tick (*Ixodes scapularis)*, a mite (*Tetranychus urticae*) and a horseshoe crab (*Limulus polyphemus*) as well as representatives of Myriapoda (*Strigamia maritima)*, Crustacea (*Daphnia pulex*) and Insecta (*Drosophila melanogaster*). Topology is based on (Sharma et al., 2014a).

As a step towards this goal, we report here the sequencing and analysis of the genomes of the common house spider *Parasteatoda tepidariorum* (C. L. Koch, 1841; formerly *Achaearanea tepidariorum*) (Yoshida, 2008) and the bark scorpion *Centruroides sculpturatus* (Wood, 1863) (Fig. 1) together with comparative genomic analyses of other available chelicerate genomes. We found that the genome of *P. tepidariorum* contains many paralogous genes including two Hox gene clusters, which is also the case in other spiders and in scorpions (this work; Di et al., 2015). These similar patterns of gene duplication between spiders and scorpions are consistent with recent molecular phylogenies, which support a much closer phylogenetic relationship of spiders and scorpions than previously thought, in a clade known collectively as Arachnopulmonata (Sharma et al., 2014a) (Fig. 1). We also document extensive divergence in the timing and location of expression of each pair of Hox gene paralogs, suggesting there may be far reaching functional consequences. Furthermore, analysis of synteny among paralogs across the *P. tepidariorum* genome is consistent with a WGD. Comparison with other chelicerates suggests that this WGD took place in the common ancestor of the Arachnopulmonata and is probably independent of the WGDs in the horseshoe crab lineage.

These results reveal that spiders and scorpions are likely the descendants of a polyploid ancestor that lived more than 450 MYA. Given the extensive morphological diversity and ecological adaptations found among these animals, rivaling those of vertebrates, our study of the ancient WGD event in Arachnopulmonata provides a new comparative platform to explore common and divergent evolutionary outcomes of polyploidization events across eukaryotes. Moreover, since *P. tepidariorum* is arguably the primary chelicerate model system in the field of evolutionary development biology (Hilbrant et al., 2012; McGregor et al., 2008; Oda and Akiyama-Oda, 2008; Schwager et al., 2015), its genome sequence will provide an excellent resource to functionally test hypotheses based on genomic inferences.

## Results

### *Parasteatoda tepidariorum* has many duplicated genes

The final *P. tepidariorum* genome assembly has a size of 1443.9 Mb. The number of predicted protein-coding genes in *P. tepidariorum* (27,990) is consistent with those of another spider S. *mimosarum* (27,235) (Sanggaard et al., 2014) as are the numbers of predicted genes of the two scorpions *M. martensii* (32,016) (Cao et al., 2013) and *C. sculpturatus* (30,456) (this study). Spiders and scorpions have significantly higher numbers of predicted genes than other arachnids such as the mite *Tetranychus urticae* (18,414) (Grbic et al., 2011). We evaluated the completeness of the *P. tepidariorum* gene set and assessed the extent of gene duplication using 429 benchmarked universal single-copy ortholog (BUSCO) groups of arthropod genes (Simao et al., 2015). For *P. tepidariorum* the HMMER3 homology search revealed 91% complete single-copy orthologs (C), 41% complete duplicated orthologs (D), and 6.5% fragmented orthologs (F). Only 2% of conserved BUSCO groups from the universal ortholog arthropods database were missing (M) from the assembly. The number of duplicated orthologs was very high compared to *Drosophila melanogaster* (C:99%, D:3.7%, F:0.2%, M:0.0%, 13,918 genes in total) or *Caenorhabditis elegans* (C:90%, D:11%, F:1.7%, M:7.5%, 20,447 genes in total).

We then undertook a different approach to further investigate the extent of gene duplication by estimating the ratios of orthologs in arachnopulmonate and non-arachnopulmonate genomes. Specifically, we compared the *P. tepidariorum* and *C. sculpturatus* genomes to the genomes of four other arthropods with a single Hox cluster and no evidence of large-scale gene duplication (“1X genomes”), including another chelicerate (the tick *Ixodes scapularis*) and three mandibulates (the red flour beetle *Tribolium castaneum*, the crustacean *Daphnia pulex*, and the centipede *Strigamia maritima*). The Orthologous Matrix (OMA) (Altenhoff et al., 2015) algorithm was used to identify orthologs after pairwise mapping of genomes. The orthology mapping indicated that, depending upon the 1X genome used for comparison, between 7.5-20.5% of spider genes that could be mapped to a single mandibulate or tick ortholog had undergone duplication (Table S1). Using the well-annotated *T. castaneum* genome as the reference, we found that 14.6% (523) of the *P. tepidariorum* genes with a single *T. castaneum* ortholog had undergone duplication (Table S1). We obtained similar results when comparing the genome of the scorpion *C. sculpturatus* with that of *T. castaneum* (10.1%, 290 genes). However, only 4.9% (175) of *I. scapularis* genes had been duplicated since its divergence from *T. castaneum* (Table S1). Moreover, higher numbers of 1:1 orthologs were found among 1X genomes than in comparisons that included either the spider or the scorpion genome, which is consistent with a greater degree of paralogy in the spider and scorpion genomes. The highest proportion of duplicated genes in a 1X genome with reference to *T. castaneum* was found in *D. pulex* (7.8%), which is known to have a large number of tandemly duplicated gene clusters (Colbourne et al., 2011) (Table S1).

Most of the spider and scorpion duplicates occurred in 1:2 paralogy (i.e., two copies in spiders/scorpions for a given mandibulate or tick homolog) (Fig. 2; Table S1), whereas duplicates in other arthropods showed no particular enrichment for this category. Two-copy duplicates accounted for 5.9-10.9% of the total spider duplicated genes, and 7.4-13.5% of the total scorpion duplicated genes (depending on the mandibulate or tick genome used for comparison). In both cases, these proportions were significantly higher than that of other arthropod genomes (*p* = 6.67 × 10^−4^) (Fig. 2A). Intriguingly, 11.8% of the 2-copy duplicates were shared between spiders and scorpions. Inversely, comparing either *P. tepidariorum* or *C. sculpturatus* to mandibulate or tick genomes recovered a much lower proportion of single copy orthologs (i.e. 1:1), relative to comparisons of any two species of mandibulate or tick. The number of duplicated genes was significantly higher in scorpions and spiders, relative to comparing mandibulate or ticks among themselves, and particularly so for the 1:2 paralog bin (two-sample *t* test; *p* = 3.75 × 10^−4^) (Fig. 2B; Table S1). We found very similar profiles of paralog distributions using a more conservative approach comparing the spider and scorpion genes to a benchmarked set of 2,806-3,031 singlecopy genes common to arthropods (the BUSCO-Ar database of the OrthoDB project) (Fig. 2C,D). Even within this database, which contained genes with no reported cases of duplication in all other studied arthropods, a considerable fraction of genes was found in two copies in both the *P. tepidariorum* and *C. sculpturatus* genomes (63-78 genes) when compared to the mandibulate or tick datasets (Fig. 2C,D; Table S1).

**Figure 2.**
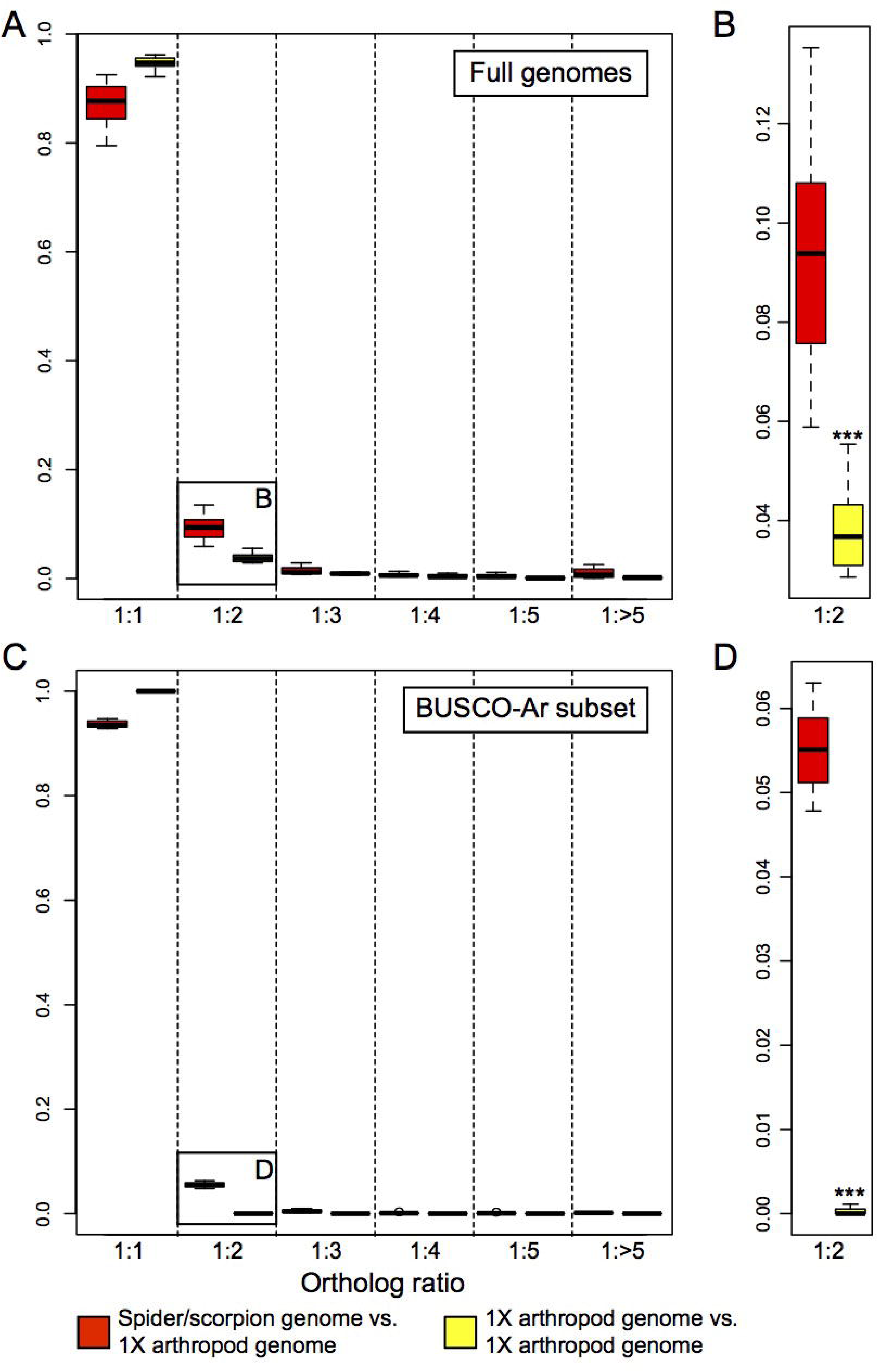
Orthology inference suggests substantial duplication in spiders and scorpions. (A) Distribution of orthology ratios from OMA analysis of full genomes. Comparisons of an arachnopulmonate genome to a 1X genome are shown in red and comparisons within 1X genomes are shown in yellow. A significantly higher number of 1:1 orthologs is recovered in pairwise comparisons within the non-arachnopulmonate genomes (p = 1.46 x 10^−3^). (B) Magnification of the 1:2 ortholog ratio category in (A) shows a significantly higher number of duplicated genes in comparisons of spider or scorpion genome to a 1X genome (p = 6.67 x 10^−4^). (C) Distribution of orthology ratios for a subset of genes benchmarked as putatively single copy across Arthropoda (BUSCO-Ar). As before, a significantly higher number of 1:1 orthologs is recovered within the 1X genome group (p = 3.43 x 10^−8^). (D) Magnification of the 1:2 ortholog ratio category in (C) shows a significantly higher number of duplicated genes in spiders and scorpions (p = 7.28 x 10^−9^).

### Dispersed and tandem gene duplicates abound in spiders and scorpions

We carried out systematic analysis of the frequency and synteny of duplicated genes in *P. tepidariorum* compared to *C. sculpturatus* and the horseshoe crab *Limulus polyphemus.* The genome of *P. tepidariorum* is characterized by an elevated number of tandem (3726 vs. 1717 and 2066 in *C. sculpturatus* and *L. polyphemus*, respectively) and proximal duplicates (2233, vs. 1114 and 97), i.e. duplicates found less or more than 10 genes away from their paralog (Figs S1-S3). However, the most salient aspect in all three genomes was the very high number of dispersed duplicates, i.e., genes for which paralogous gene models were detected on different scaffolds, amounting to approximately 14,700 genes in each species (Figs S1-S3).

To better understand the patterns of gene duplication in *P. tepidariorum*, we next investigated the duplication level and colinearity of specific coding and non-coding genes. We identified 80 homeobox gene families in *P. tepidariorum* (Table S2) of which 59% were duplicated, with a total of 146 genes (Fig. 3). Note that a very similar repertoire was also observed in *C. sculpturatus* where 60% of homeobox gene families were duplicated (156 genes representing 82 gene families (Table S3)). Of the 47 and 49 homeobox gene families with multiple gene copies in *P. tepidariorum* and *C. sculpturatus* respectively, 39 were common to both species. In addition, 22 families were represented by a single gene in both the spider and scorpion genomes (Fig. 3). The few remaining families contained duplicates in only one of these two species or were only found in one species (Fig. 3). In addition, one family, *Dmbx*, had two copies in *P. tepidariorum* but was missing in *C. sculpturatus.*

**Figure 3.**
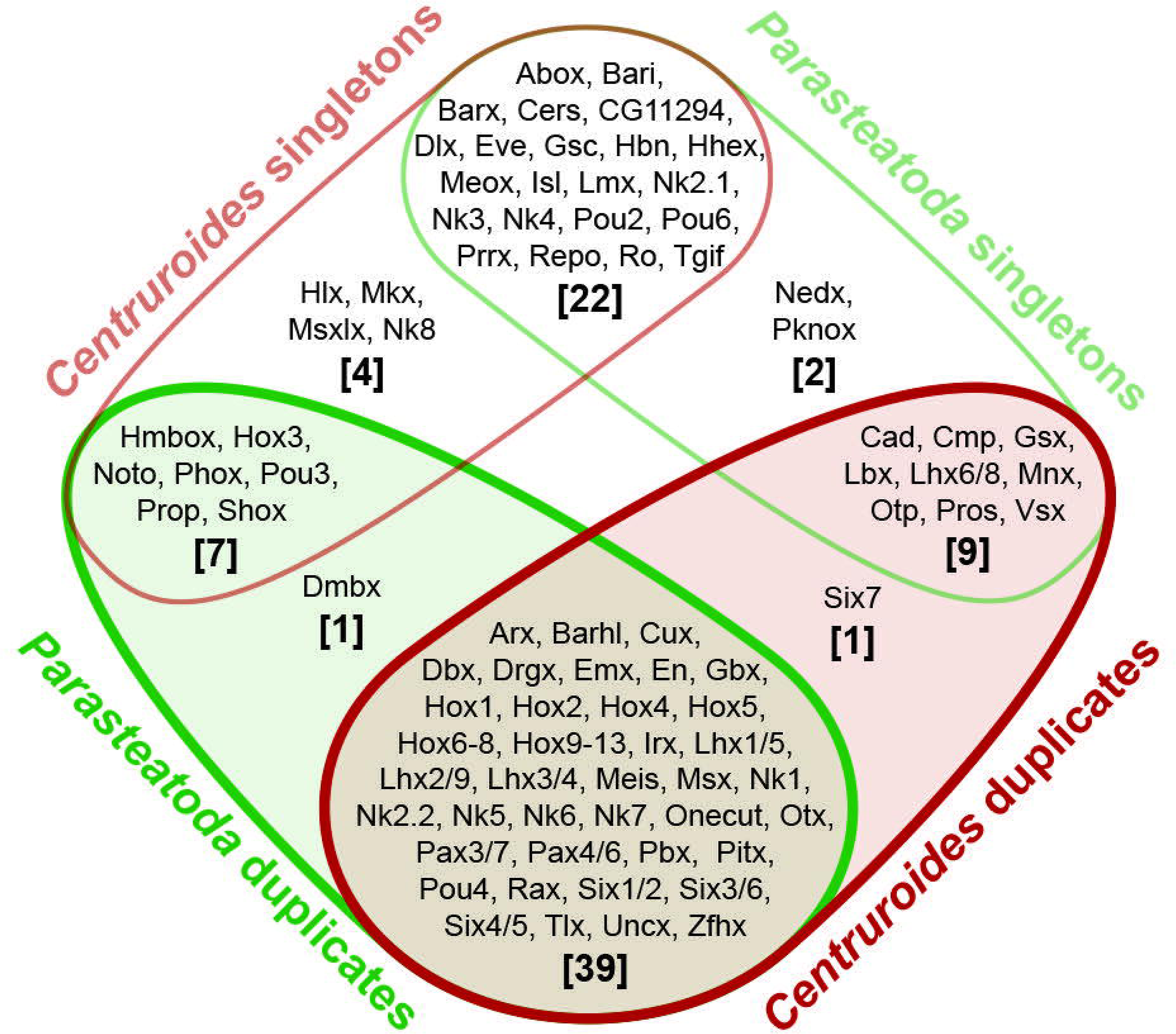
Homeodomain containing genes are frequently duplicated in *P. tepidariorum* and *C. sculpturatus*. Many duplicated homeobox gene families (overlap of red and green shading) are shared between *P. tepidariorum* (indicated in green) and *C. sculpturatus* (indicated in red). Single copy families are the next largest group shared, then families that are single copy in one species but duplicated in the other. There are also a few families that were only found in one species.

The duplication of Hox gene clusters in vertebrates was among the first clues that led to the discovery of ancient WGDs in this group (Amores et al., 1998). Therefore we assessed the repertoire and organization of Hox genes in *P. tepidariorum* in comparison to three other spider genomes (*L. hesperus, S. mimosarum* and *A. geniculata*; Sanggaard et al. (2014)), two scorpion genomes (C. *sculpturatus* and *M. martensii*; Cao et al. (2013), this study) and the tick genome (*I. scapularis*; Gulia-Nuss et al. (2016); Lawson et al. (2009)).

We identified and manually annotated orthologs of all ten arthropod Hox gene classes (*labial* (*lab*), *proboscipedia* (*pb*), *Hox3, Deformed* (*Dfd*), *Sex combs reduced* (Scr), *fushi tarazu* (*ftz*), *Antennapedia* (*Antp*), *Ultrabithorax* (*Ubx*), *abdominal-A* (*abdA*) and *Abdominal-B* (*AbdB*)) in all genomes surveyed (Fig. 4; Table S4). Whereas the tick genome contains only one copy of each Hox gene, nearly all Hox genes are found in two copies in the spider and scorpion genomes (Fig. 4; Fig. S4; Table S4). The only Hox gene is not always found in duplicate is *ftz* in *P. tepidariorum* (see Fig. 4; Fig. S4; Table S4).

**Figure 4.**
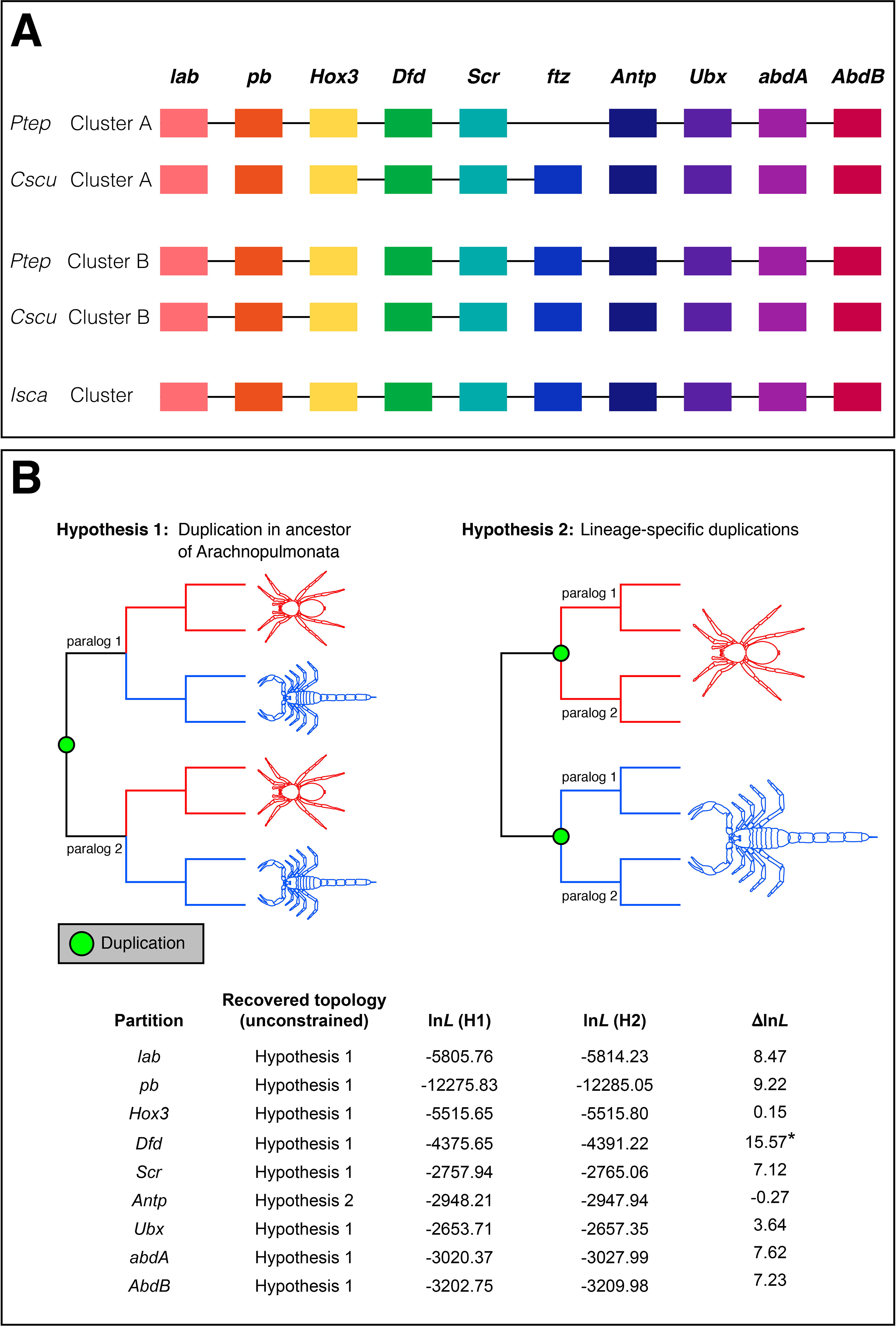
Hox gene complement and hypothetical Hox clusters in chelicerate genomes. Hox gene clusters A and B in the spider *Parasteatoda tepidariorum* and the scorpion *Centruroides sculpturatus*, and in the tick (A). For details, see Table S4. Transcription for all genes is in the reverse direction. Genes (or fragments thereof, see Table S4) that are found on the same scaffold are joined by black horizontal lines. Abbreviations: *Ptep Parasteatoda tepidariorum, Cscu Centruroides sculpturatus, Isca Ixodes scapularis.* (B) Gene tree analysis of individual Hox genes support a shared duplication event in the common ancestor of spiders and scorpions in all cases except *Antennapedia.*

Interestingly, none of the Hox paralogs present in spiders and scorpions were found as tandem duplicates. Instead, in *P. tepidariorum*, the species with the most complete assembly in this genomic region, it was clear that the entire Hox cluster had been duplicated *in toto*, We found one *P. tepidariorum* Hox cluster copy in a single scaffold, lacking only a *ftz* copy, which this particular cluster is lacking presumably in all spiders (Fig. 4, Table S4). The second Hox cluster was split between two scaffolds, which could be due to the incomplete assembly of this region because there is not enough sequence downstream of *Dfd* (~70 kb) and upstream of Hox3 (~320 kb) to cover the paralogous ~840 kb between *Dfd* and *Hox3* on Cluster A in *P. tepidariorum* or even the ~490 kb between *Dfd* and *Hox3* in *I. scapularis* (Table S4).

Comparative analyses suggests that other key developmental genes are also commonly duplicated in *P. tepidariorum.* A synteny analysis of these previously reported duplications showed that only the two *Pax6* paralogs were located on the same scaffold (Table S5), suggesting that they arose through tandem duplication. The paralogs of other duplicated developmental genes examined were found on different scaffolds (Table S5): retinal differentiation (*dachshund* and *sine oculis*), head patterning (*six3, orthodenticle, collier*) (Samadi et al., 2015; Schomburg et al., 2015), Wnt pathway genes (*Wnt7, Wnt11, frizzled 4*) (Janssen et al., 2010; Janssen et al., 2015) and appendage formation genes (*homothorax, extradenticle, Lim1, spineless, trachealess* and *clawless*) (Prpic et al., unpublished data).

It was also recently reported that approximately 36% of annotated microRNAs in *P. tepidariorum* are present as two or more copies (Leite et al., 2016). Analysis of the synteny of the paralogous *P. tepidariorum* microRNAs shows that only 8 out of 30 are found on the same scaffold. Furthermore, nearly all of the tandemly duplicated microRNAs in *P. tepidariorum* are microRNAs that are largely specific to this spider (e.g. mir-3971 paralogs) or are clustered in arthropods (e.g. mir-2 from the mir-71/mir-2 cluster) (Table S6) (Leite et al., 2016). These findings suggest that the majority of duplicated microRNAs were not generated by tandem duplication.

Taken together, classification of duplicated genes in spiders and scorpions show that tandem and especially dispersed duplications abound in these genomes. Furthermore, a Hox cluster duplication could be indicative of a WGD.

### Conservation of synteny among *P. tepidariorum* scaffolds supports the hypothesis of a WGD event

To further test the hypothesis that a WGD event had occurred in an ancestor of *P. tepidariorum*, we next searched for conserved synteny among the genomic scaffolds of this spider using Satsuma (Grabherr et al., 2010) (note that this approach was not possible in *C. sculpturatus* because of assembly quality of the genome of this scorpion). This analysis revealed signatures of large segmental duplications suggestive of a WGD followed by numerous rearrangements (inversions, translocations, tandem duplications) (Fig. 5A). These signatures were observed among many of the larger scaffolds (Figs 5 and S5), but were particularly strong and clear between scaffolds 1 and 7, between scaffolds 9 and 30, and among scaffolds 60, 78 and 103 (Fig. 5B). These results are similar to findings from a similar analysis of the genome of the fish *Tetraodon nigroviridis* (Jaillon et al., 2004) and are consistent with an ancient WGD event in an ancestor of this spider.

**Figure 5.**
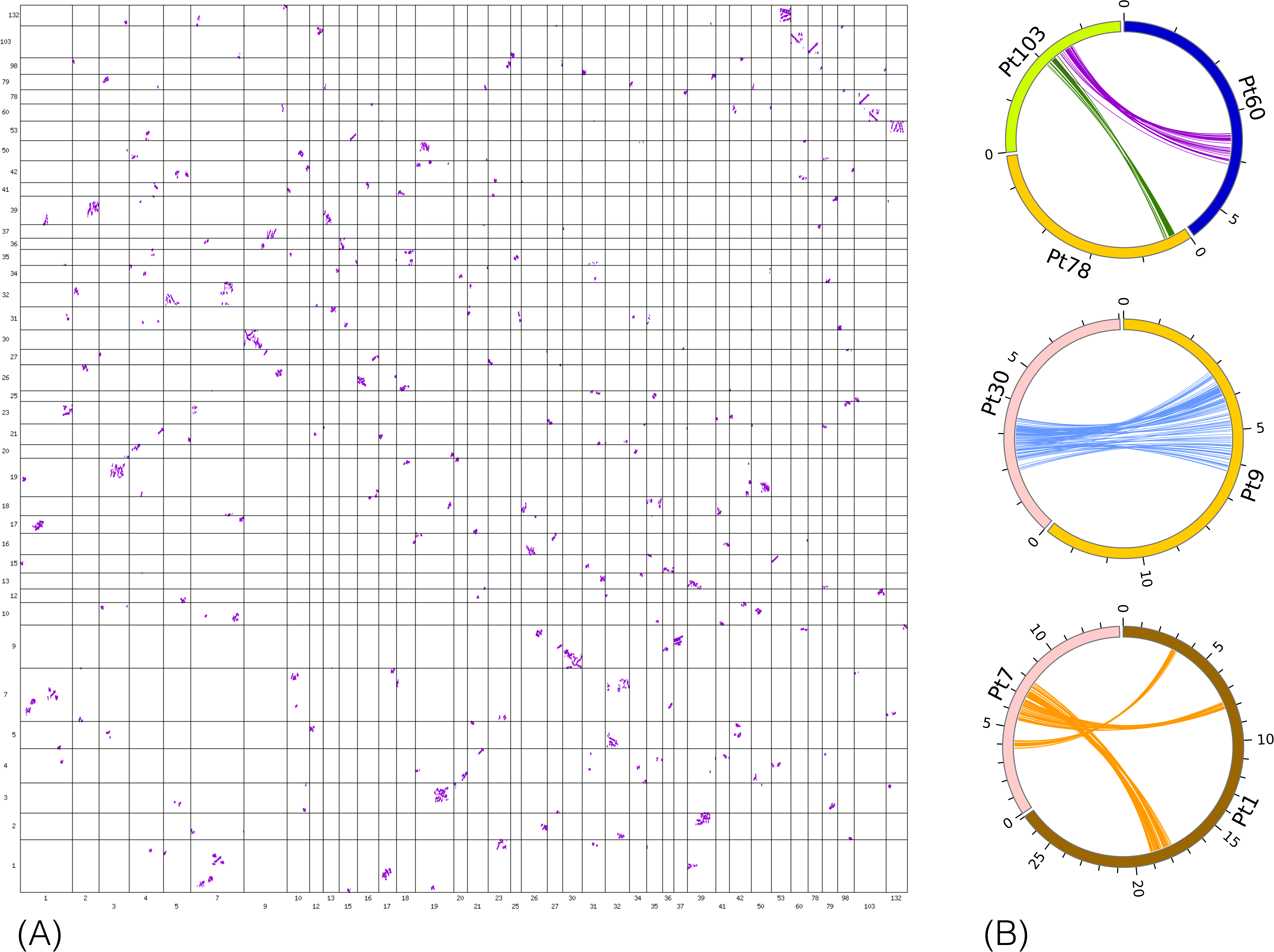
Genome-scale conservation of synteny among *P. tepidariorum* scaffolds reveals signatures of an ancient WGD. (A) Oxford grid displaying the colinearity detected by SatsumaSynteny among the 39 scaffolds presenting the greatest numbers of hits on one another. On this grid (not drawn to scale), each point represents a pair of identical or nearly identical 4096 bp region. Alignments of points reveal large segmental duplications suggestive of a whole-genome duplication even, along with other rearrangements such as inversions, translocations and tandem duplications. (B) Circos close-ups of some of the colinearity relationships revealed by the Oxford grid.

### When did WGD occur in chelicerates?

To determine the timing of duplication relative to species divergence within a broad taxonomic sampling of arachnids than analyzed thus far, we grouped the protein-coding genes of 30 arachnid species into gene families with either *P. tepidariorum* or *C. sculpturatus* translated genes used as a seed plus *L. polyphemus* and S. *maritima* as outgroups (Table S7) (Goodstein et al., 2012). This method resulted in 2,734 unique *P. tepidariorum-seeded* gene families (Fig. S6). Note that seeding gene families with *C. sculpturatus* resulted in fewer families (1,777) but similar patterns of gene duplication (not shown); we thus focused on the results of *P. tepidariorum-seeded* families.

To analyze the timing of the putative WGD event, we calculated molecular distances between paralog pairs by averaging the maximum likelihood branch lengths estimated under the HKY model of evolution (Hasegawa et al., 1985) within gene trees from the duplication node to all descendant within-species paralogs. We fit the molecular distances of duplication nodes with HKY > 0.01 (avoid inferring alleles as paralogs) and HKY < 2.0 (minimize mutational saturation) to five distribution models. The results show that *P. tepidariorum* duplication nodes best fit three Gaussian distributions (four other distributions were rejected by the Kolmogorov-Smirnoff Goodness-of-Fit test, Table S8. The first Gaussian distribution, with an average genetic distance of μ = 0.038 likely represents recent individual gene duplications. The second (μ = 0.491) and third (μ = 1.301) distributions of genetic distance among paralogs are consistent with two ancient large-scale duplication events (Fig. 6A) (Kagale et al., 2014; Nossa et al., 2014). We observed a similar distribution of paralog molecular distances in five deeply sequenced spider species and *C. sculpturatus* (Table S9, Fig. S7), but not *T. urticae* and *I. scapularis.* The shift in distribution patterns between the scorpion and the mite is consistent with a shared WGD in spiders and scorpions that is not experienced by the more distantly related arachnid species. It is also possible that spiders and scorpions experienced independent duplication events shortly after their divergence, but this is unlikely given the shared retention of paralogs from this analysis (see below) and from the BUSCO-Ar and OMA gene sets (see above).

**Figure 6.**
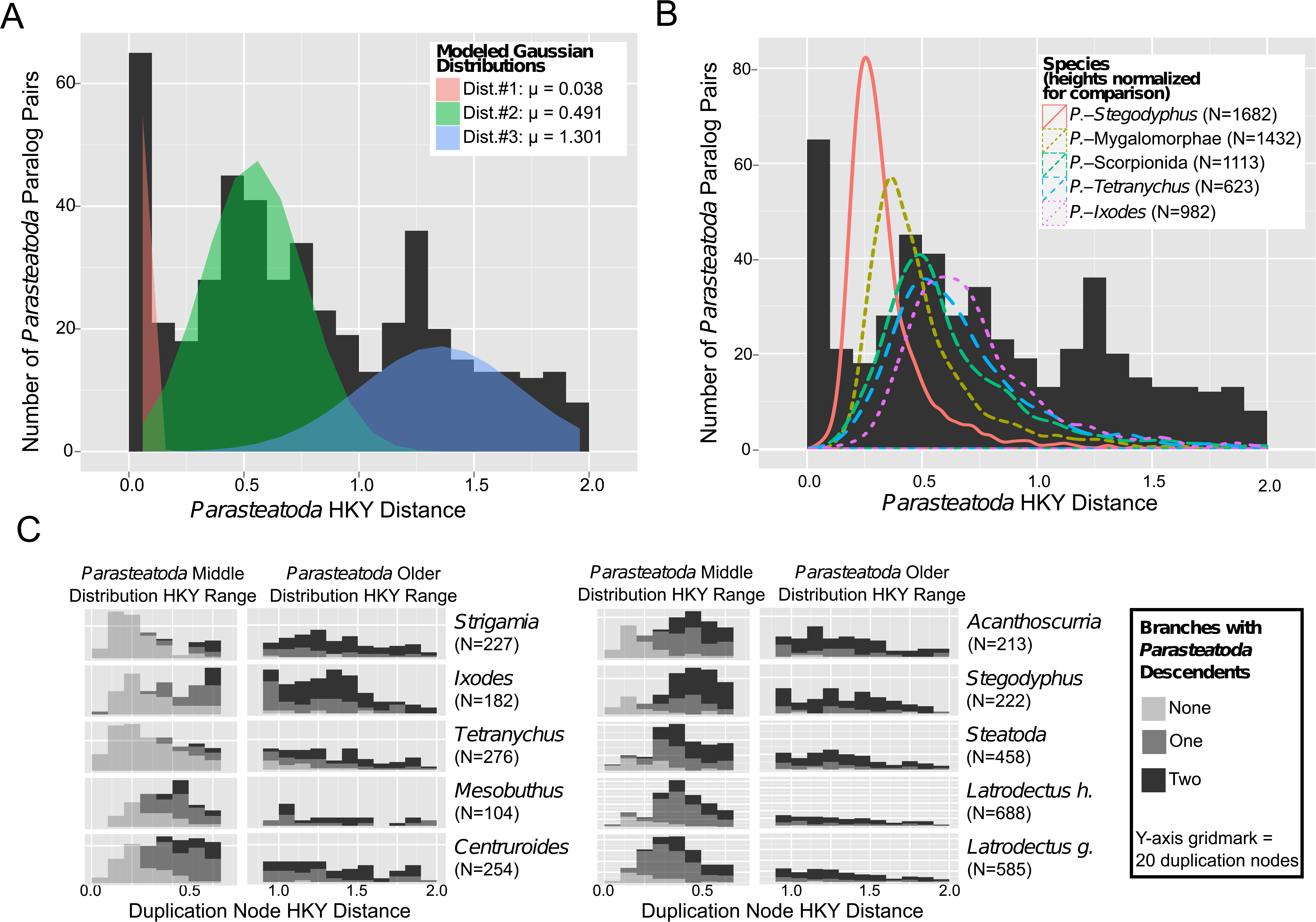
Molecular Distance Distributions of *P. tepidariorum* Paralogs and Speciation Nodes. The distribution of mean HKY distances from *P. tepidariorum* duplication nodes to *P. tepidariorum* descendants reveals three distributions shown in different colors in (A). Comparing the distribution of HKY distances from speciation nodes to *P. tepidariorum* (lines in B) reveals that distribution #1 (red in A) is restricted to the *P. tepidariorum* branch, distribution #2 (green in A) is similar to pre-spider and post-tick speciation nodes, and distribution #3 (blue in A) is older than the *P. tepidariorum*-tick speciation event. N=number of speciation nodes in B. Comparing the number of duplication nodes in non-*P. tepidariorum* species (C) that are either partially or fully retained in *P. tepidariorum* reveals the duplication nodes with HYK distances in the range of the oldest *P. tepidariorum* distribution (blue in A) are retained at a similar rate across all species (right subcolumns in C), but that those duplication nodes with HKY distances in the range of the middle *P. tepidariorum* distribution (green in A) are only retained in scorpions or more closely related species (left sub-columns C).

The possibility that a WGD occurred prior to the divergence of spiders and scorpions and after the divergence of spiders from mites is additionally supported by comparison of the distributions of HKY distances of the duplication nodes to speciation nodes, with an almost identical pattern found for the paralog distances and the spider-scorpion distances (Figs 6B, S8, Table S10). Shared paralog retention is also high for spiders and scorpions, but not between spiders and ticks or mites, further supporting a shared WGD in the spider and scorpion common ancestor (Fig. 6C; Table S11). Furthermore, tandem duplication nodes identified above formed a majority of the duplication nodes in the younger Gaussian distribution (71%), and minorities of the second (24%) and third distributions (9%) (Fig. S9). This is the opposite of what is seen with the duplication nodes containing dispersed duplications (younger: 29%, second: 62%, and third: 50%). Additionally, a slight majority of the older tandem duplication nodes showed evidence of being shared with other arachnids (57%), but mostly with other species in the same family as *P. tepidariorum* (44%). This suggests that an ancient WGD was followed by pervasive lineage specific tandem duplications especially in spiders.

Analysis of the gene families containing a duplication pair from the middle and oldest Gaussian distributions (Fig. 6A), excluding tandem duplicates, showed that they are enriched in several GO terms compared to gene families without duplication pairs, including several terms associated with transcription, and metabolism (Table S12). The same GO terms are also enriched in these gene families compared to the families with tandem duplications, but the difference is not significant. However, the gene families with tandem duplication pairs are depleted in GO terms relating to translation.

### Gene trees support the common duplication of genes in Arachnopulmonata

The results of our analysis of duplicated genes in *P. tepidariorum* and other arachnids from the OMA and BUSCO gene sets, as well as our dating of the divergence in gene families, strongly suggest that there was a WGD in the ancestor of spider and scorpions. To further explore whether the duplicated genes in spiders and scorpions were the result of duplication in the most recent common ancestor of these arachnopulmonates (Hypothesis 1) or lineage specific duplications (Hypothesis 2), we applied a phylogenetic approach to examine *P. tepidariorum* and *C. sculpturatus* genes (Fig. 7; Tables S13, S14). Of the 116 informative gene trees (see methods) of orthogroups wherein exactly two *P. tepidariorum* paralogs were present for a single *T. castaneum* ortholog, 67 (58%) (henceforth Tree Set 1) were consistent with a common duplication (Hypothesis 1) and 49 (42%) were consistent with lineage specific duplications (Hypothesis 2) (Fig. 7; Tables S13, S14). Of the 67 tree topologies supporting a common duplication, 18 were fully congruent with the idealized Hypothesis 1 tree topology, and 49 were partially congruent with Hypothesis 1 (i.e., the two spider paralogs formed a clade with respect to a single scorpion ortholog) (Fig. 7; Tables S13, S14).

**Figure 7.**
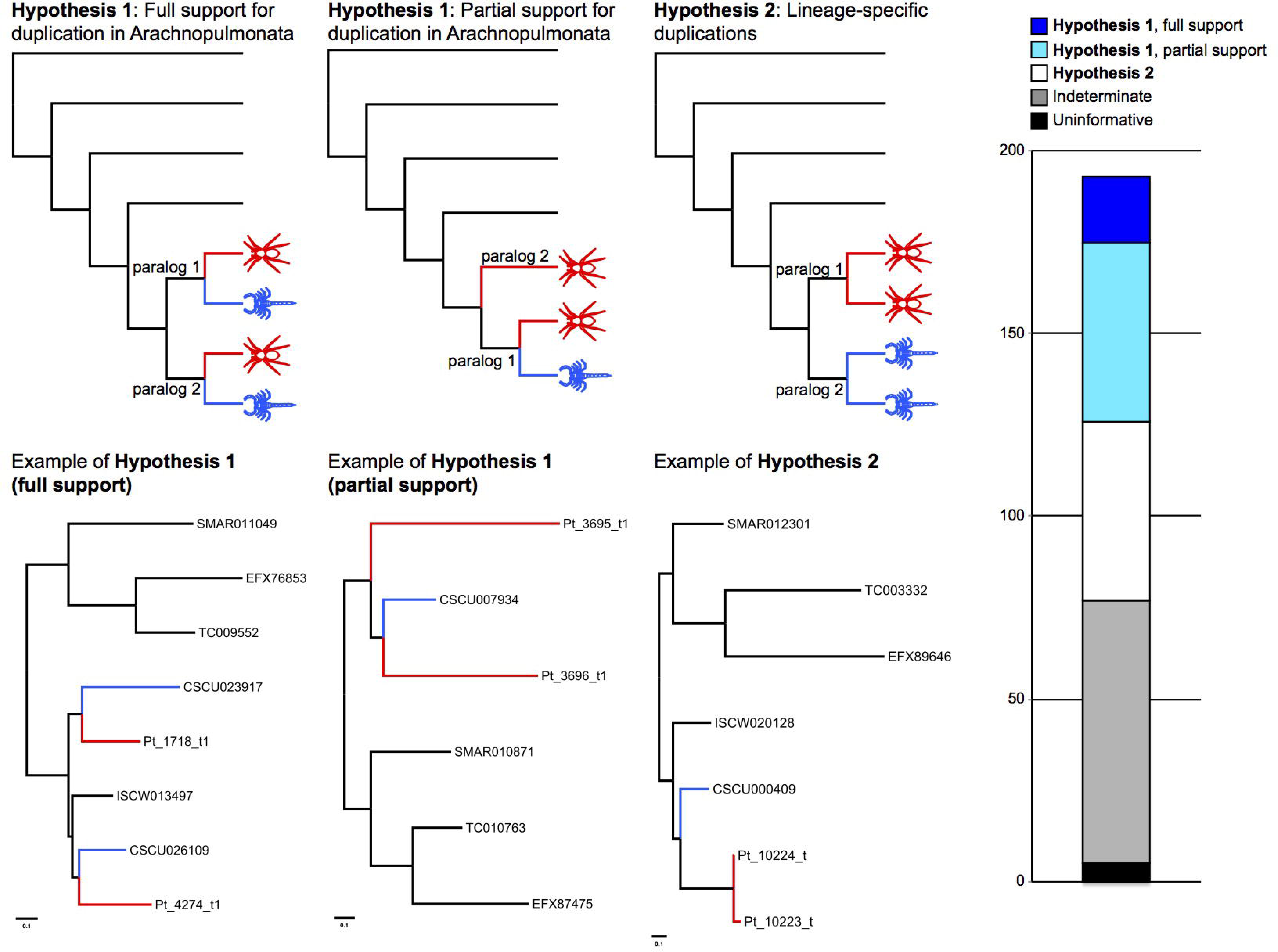
Gene trees support the common duplication of genes in Arachnopulmonata. Analysis of gene trees inferred from six arthropod genomes was conducted, with the gene trees binned by topology. Trees corresponding to a shared duplication event were binned as Hypothesis 1, and trees corresponding to lineage-specific duplication events as Hypothesis 2. Gene trees with spider paralogs forming a grade with respect to a single scorpion paralog were treated as partially consistent with Hypothesis 1. Top row of panels shows hypothetical tree topologies; bottom row of panels shows empirical examples. Right panel shows distribution of gene trees as a function of bin frequency.

If the gene trees in Tree Set 1 were the result of large-scale duplication events or WGD as opposed to tandem duplication, we would expect each resulting copy to occupy two different scaffolds. Of the 18 *P. tepidariorum* paralog pairs from gene trees fully consistent with Hypothesis 1, 15 were found to occupy different *P. tepidariorum* scaffolds; of the 49 paralogs pairs from gene trees partially congruent with Hypothesis 1, all but ten pairs were found to occupy different *P. tepidariorum* scaffolds (Table S15). In addition, of the 18 *C. sculpturatus* paralog pairs that were fully consistent with Hypothesis 1, all 18 were found on different scaffolds. To test whether *P. tepidariorum* paralog pairs inferred to inhabit different scaffolds were distinguishable in assembly completeness from the three paralog pairs on the same *P. tepidariorum* scaffolds, we examined the length of the scaffolds for these two groups. We found the lengths of the scaffolds were statistically indistinguishable between the two groups (Table S15; Wilcoxon rank sum test: W = 358, p-value = 0.9179). This analysis was not required for the 18 scorpion paralogs pairs, because in all cases, each member of the scorpion paralog pair was distributed on a different scaffold.

The occurrence of two clusters of Hox genes in both the spider and scorpion genomes could also be consistent with either of these alternative hypotheses (Fig. 4B). However, only in the case of *Antp* was tree topology consistent with Hypothesis 2 recovered and the difference in log likelihood with respect to the log likelihood topology was negligible (ln*L* = −0.27) (Fig. 4B). Higher statistical support for the Hypothesis 1 topology was generally obtained for data partitions with a large number of available sequences (e.g., *Dfd, pb*) (Fig. 4B). The sum of the Hox gene tree data is therefore consistent with the synteny analysis, and supports a shared duplication in the common ancestor of Arachnopulmonata.

### WGD in Xiphosura is probably unrelated to the duplication of genes in Arachnopulmonata

The recent report of WGD and multiple Hox clusters in an analysis of horseshoe crabs (Order Xiphosura; Kenny et al. (2016)) raises the possibility of two alternative interpretations: (a) a single WGD at the base of Chelicerata, with losses of duplicated genes in lineages like mites and ticks, or (b) separate WGD events in the horseshoe crab ancestor and in the arachnopulmonate ancestor. To discern whether the WGD event/s recently reported in Xiphosura constitute separate (Hypothesis 3) or common (Hypothesis 4) evolutionary events from the duplication of genes in Arachnopulmonata, we added the three published horseshoe crab genomes to our dataset and reran OMA (Fig. 8). If the duplications reported here in spiders and scorpions were caused by the same event that drove the genome duplications in horseshoe crabs, we would expect to find paralog clusters that included members of all Euchelicerata (Xiphosura + Arachnida). This expected pattern is comparable to the case of whole genome duplications in the vertebrate ancestor (Dehal and Boore, 2005), which resulted in the same sets of paralogs for all major vertebrate lineages, to the exclusion of non-vertebrate deuterostomes and the protostomes (e.g., the *Sp* gene family; Schaeper et al. (2010)). By contrast, if the duplications in spiders and scorpions were distinct from the duplications in horseshoe crabs, we would expect to observe a pattern where (a) horseshoe crab paralogs clustered together, (b) arachnopulmonate paralogs clustered together, and (c) all other arachnid orthologs would not be duplicated at all, and fall somewhere in between horseshoe crabs and arachnopulmonates (Fig. 1; Sharma et al. (2014a)). We thus examined gene trees recovered by OMA to discern which of these two scenarios was supported by the comparison of the nine full genomes.

**Figure 8.**
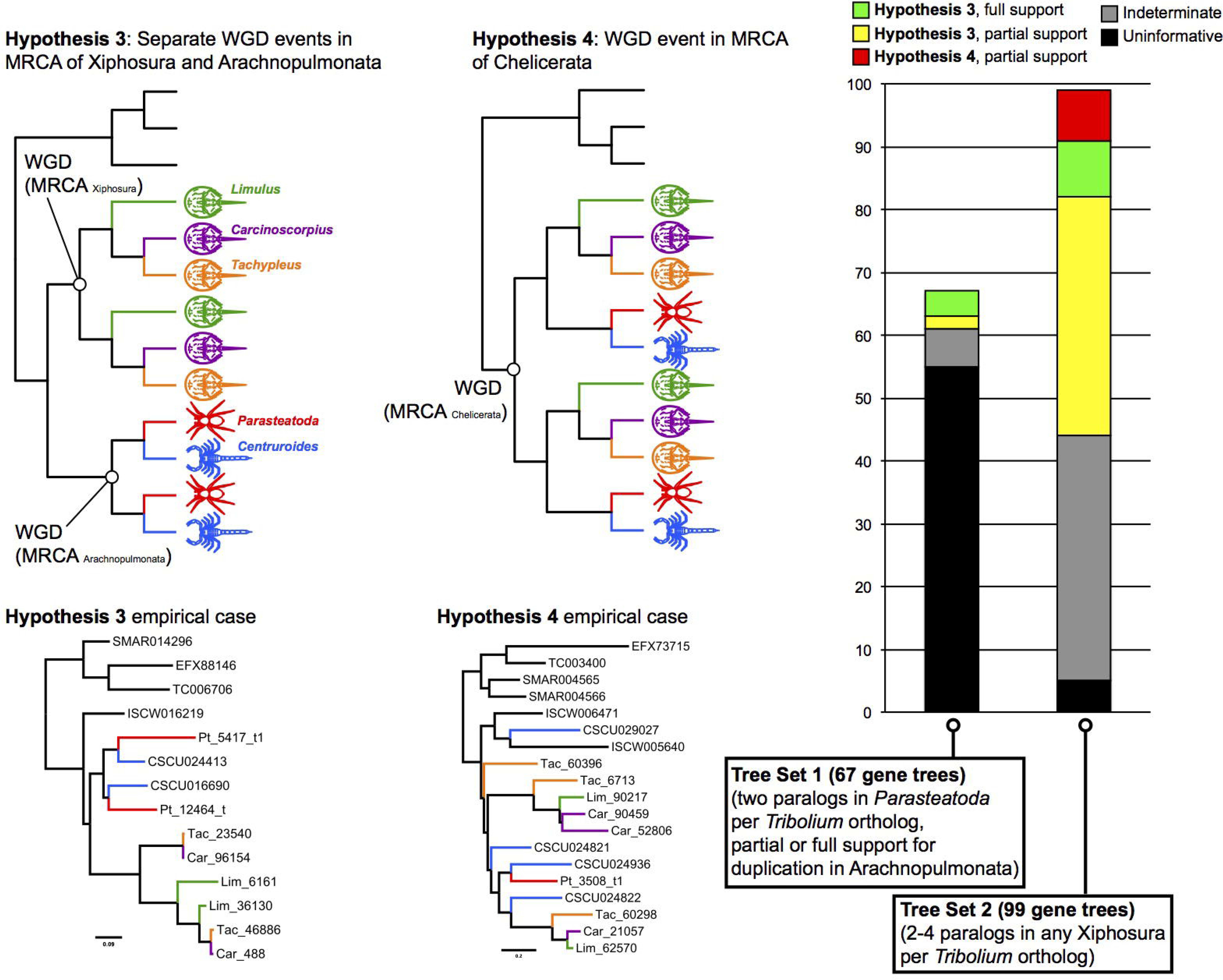
WGD in Xiphosura is probably unrelated to the duplication of genes in Arachnopulmonata. Analysis of gene trees inferred from nine arthropod genomes was conducted, with the gene trees binned by topology. Trees corresponding to two separate duplication events in the MRCA of Xiphosura and Arachnopulmonata were binned as Hypothesis 3, and trees corresponding to a single duplication events in the MRCA of Chelicerata as Hypothesis 4. Top row of panels shows hypothetical tree topologies; bottom row of panels shows empirical examples. Right panel shows distribution of gene trees as a function of bin frequency, for two different tree sets (i.e., gene trees retrieved under two alternate filtering criteria). Note the limited support for Hypothesis 4, with empirical gene trees poorly matching the expected tree topology (contra empirical cases supporting Hypothesis 3).

We first examined the orthogroups corresponding to Tree Set 1, after addition of horseshoe crab orthologs (Fig. 8). However, we found that 55 of the 67 gene trees constituting Tree Set 1 could not distinguish between Hypothesis 3 and Hypothesis 4 (i.e., no horseshoe crab paralogs were recovered in those orthogroups with duplicated spider genes).

We assembled a second tree set (henceforth, Tree Set 2) using the filtering criterion of orthogroups where 2-4 xiphosuran paralogs were recovered for a single *T. castaneum* ortholog. We thus recovered 99 gene trees in Tree Set 2 (Fig. 8). Of these, 44 were indeterminate (non-monophyletic outgroup) or uninformative (either missing all arachnopulmonates or missing all xiphosuran paralogs). A further 47 were consistent with Hypothesis 3, with nine gene trees completely congruent with Hypothesis 3 (i.e., multiple paralog clusters within both arachnopulmonates and horseshoe crabs; monophyly of Arachnopulmonata and Xiphosura; and monophyly of the mandibulate outgroup) (Fig. 8). The last eight gene trees in Tree Set 2 were scored as partially consistent with Hypothesis 4, but as shown in one empirical case (Fig. 8), these gene trees did not correspond well to the scenario of a common WGD at the base of Chelicerata, and may stem from algorithmic error in phylogenetic reconstruction (e.g., model misspecification). To be conservative, we treated these eight trees as consistent with our alternative hypothesis.

The sum of our gene tree analyses thus indicates support for Hypothesis 3 - the independent origins of arachnopulmonate and xiphosuran duplications. We found very little support for a shared duplication event at the base of Chelicerata (Hypothesis 4); no gene tree could be found where multiple paralogous groups each included exemplars of Xiphosura and Arachnopulmonata. Taken together, these results suggest that the duplication of genes in spiders and scorpions was probably independent of the proposed WGD events in horseshoe crabs.

### Hox gene paralogs in *P. tepidariorum* show considerable divergence in temporal and spatial expression during embryogenesis

Alteration of the temporal and/or spatial expression can underlie the neo- or sub-functionalization of duplicated genes. To test whether the Hox gene paralogs in chelicerates have divergent expression patterns, we assayed the expression of all 19 Hox genes throughout *P. tepidariorum* embryogenesis. For each pair of Hox paralogs, we found remarkable differences in spatial and temporal expression patterns (Fig. 9, Figs S10-25).

**Figure 9.**
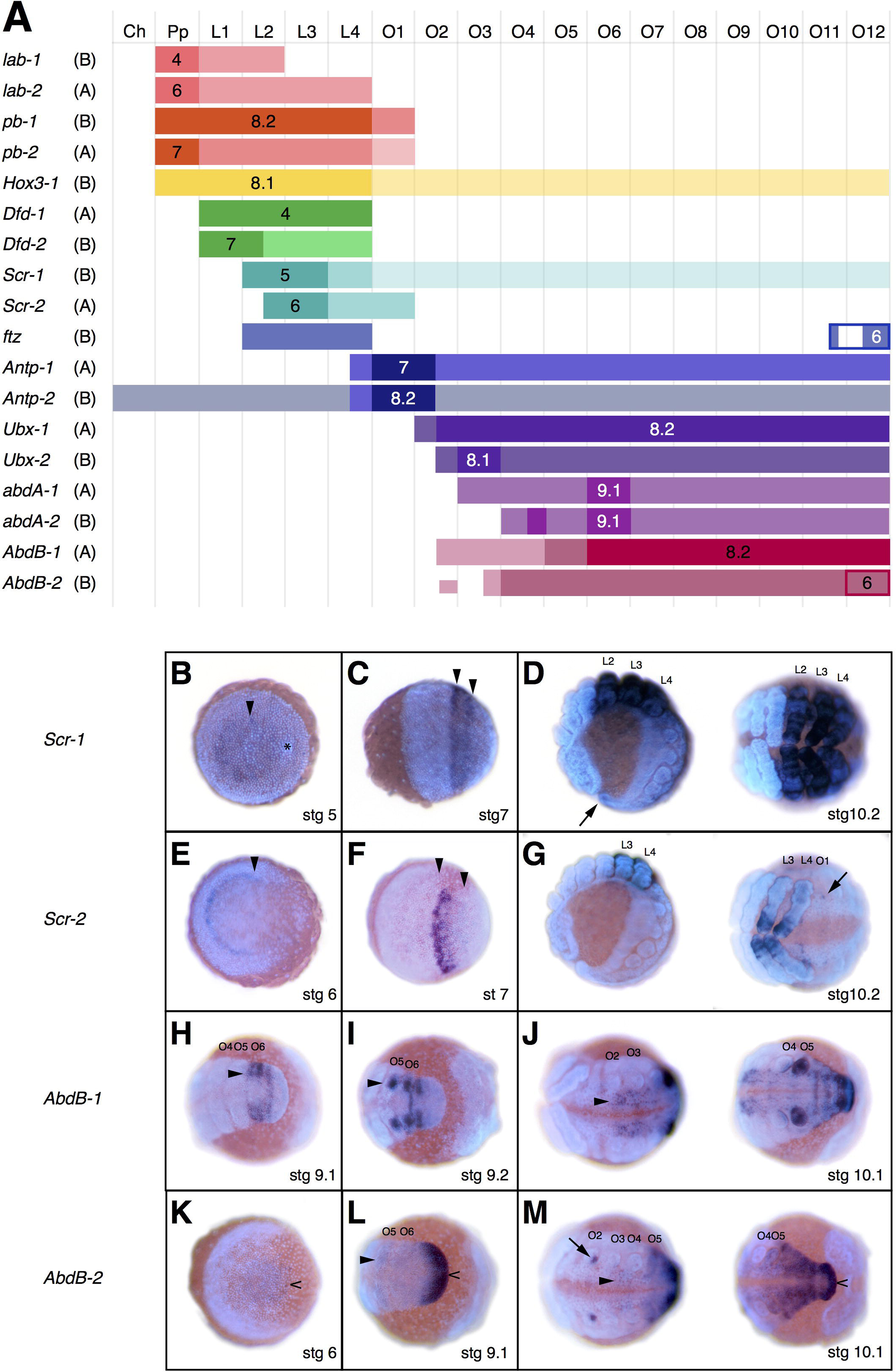
Expression of Hox paralogs in *P. tepidariorum*. (A) Summary of Hox gene expression domains and expression timing in *P. tepidariorum* embryos. Columns represent segments from anterior to posterior. Bars represent the extent of a gene’s expression domain with respect to the segments. The darkest color for each gene is used for the initial expression domain of each gene when it first appears, which usually coincides with a genes’ strongest expression. The next lighter color is used for the expanded domain, and the lightest color is used for further late expansions of the expression domains, which usually tends to be only in the nervous system. The stage at which a genes’ expression first appears, is depicted by the stage number in the domain of first expression. The affiliation to cluster A or B is noted in brackets after the gene name. *ftz*, in addition to its Hox domain is, expressed dynamically (i.e. budding off stripes) in the SAZ, and *AbdB-2* is continuously expressed in the SAZ after its formation at stage 6. These SAZ expression patterns are indicated by rectangular outlines in what is otherwise the O12 segment. Note that we did not detect any expression for *Hox3-2.* (B-M) Two examples of Hox gene expression differences between paralogs of *Scr* (B-G) and *AbdB-2* (H-M). All images are overlays of a brightfield image depicting the expression pattern and a fluorescent DAPI nuclear staining. (B) *Scr-1* is first expressed at stage 5 in a cap-like domain (arrowhead) in the center of the germ disc. The cumulus is marked by an asterisk. (C) During germ band formation, the *Scr-1* expression forms two distinct stripes in the presumptive anterior L2 and L3 segments (arrowheads), which are followed by weaker expression in the posterior segment. (D) At later stages, *Scr-1* expression (lateral view on the left, ventral view of leg region on the right) refines into a pattern consisting of multiple rings in the legs developing on L2-L4, but the expression level is always higher in L3 compared to L2 and L4. Additionally, weaker expression can be found all throughout the opisthosoma extending into the most posterior segments (arrow). (E) *Scr-2* is first expressed only at stage 6, during very early germ band formation. It is expressed in one stripe (arrowhead) that later at stage 7 (F) is located in the presumptive posterior of L2 and in L3, whereas weaker expression is found in L4 (borders marked by arrowheads). At later stages (G), expression is predominantly seen in rings L3 and L4 that are markedly different from the pattern seen for *Scr-1.* Additional expression can be seen in a few cells of the posterior half of each segments’ neuroectoderm of L2-L4 and in O1 (arrow). (H) *AbdB-1* expression is first visible at stage 9.1, and its anterior border coincides with the anterior border of O6 (arrowhead). It then expands anteriorly into the limb buds of O5 at stage 9.2 (arrowhead), and posterior segments also will express *AbdB-1* as they are added from the SAZ. (I) At later stages (J) (ventral view of the anterior opisthosoma on the left, posterior opisthosoma on the right), expression expands anteriorly into the ventral neuroectoderm of the posterior O2 segment. *AbdB-2* however, is starting to be expressed in the SAZ (caret) during its formation at stage 6 (K). It continues to be expressed strongly in the SAZ during germ band formation (caret), and the expression only expands into O5 at stage 9.1 (arrowhead) (L). (M) At later stages, *AbdB-2* also extends into the ventral neuroectoderm of O4 (arrowhead), and expression additionally appears in the genital opening on O2 (arrow), while expression continues in the SAZ (caret). Abbreviations: Ch cheliceral segment, Pp Pedipalpal segment, L-L4 walking leg segments 1-4, O1-12 opisthosomal segments 1-12.

The expression of the paralogs of each Hox gene never appears at the same time during development; the expression of one paralog often precedes the other paralog by at least 10 hours (e.g. *lab, Scr, Ubx* and *AbdA*) (Akiyama-Oda and Oda, 2010; Pechmann et al., 2015) (Fig. 9B-G), if not 15 to 20 hours (pb, *Dfd, Antp)*, and even 30 hours in the case of *AbdB* (Fig 9A, H-M). The expression domains of paralogs also differ significantly in their anterior and/or posterior borders. *Scr, Ubx, abdA* and *AbdB* paralogs exhibit anterior borders that are shifted by half a segment or more, and most prosomally expressed Hox gene paralogs show shifts in their posterior expression borders by one or more segments (Fig 9A). While the borders of the strongest expression domain are identical in the case of the paralogs of *lab, Antp*, and *abdA*, they differ substantially in all other paralogs (Fig. 9, Figs S10-25).

Most Hox gene paralogs also exhibit differences in the tissues and cells types they are expressed in (e.g. mesodermal vs. ectodermal expression, or groups of neuroectodermal cells that a paralog is expressed in), which hints at the possible neofunctionalization of one of the paralogs. For example in the case of the *AbdB* paralogs (Fig. 9H-M), only *AbdB-2*, is expressed in the segment addition zone where it has a dynamic anterior expression border until a more Hox-like expression domain appears at stage 9. Taken together, we have observed considerable differences in the spatial and temporal expression between each of the *P. tepidariorum* Hox gene paralogs (Fig. 9). These differences reflect the considerable divergence time since the Hox gene cluster was duplicated and likely also evidence changes in function between the paralogs.

## Discussion

### Signatures of an ancient WGD in the last common ancestor of spiders and scorpions

Our study of the assembly and annotation of the *P. tepidariorum* genome revealed a high number of duplicated genes in accordance with previous observations (Clarke et al., 2015; Clarke et al., 2014; Di et al., 2015; Fuzita et al., 2015; Fuzita et al., 2016; Janssen et al., 2010; Leite et al., 2016; Schwager et al., 2007; Sharma et al., 2015; Sharma et al., 2014b; Turetzek et al., 2016). This finding is further supported by our detection of a colinearity signal across many of the largest *P. tepidariorum* scaffolds. The fact that we find many smaller synteny blocks across scaffolds suggests that the WGD event occurred early during spider evolution and was followed by extensive disruption of previously larger blocks for instance by recombination or the activity of transposable elements. Intriguingly, the comparison of the gene content of the *P. tepidariorum* genome with other chelicerates and other arthropods suggests that a WGD likely occurred in the lineage leading to spiders and scorpions. Our dating efforts indeed confirmed that this WGD most likely occurred after the divergence of the common ancestor of spiders and scorpions from other arachnid lineages (mites, ticks and harvestmen) prior to 430 MYA (Dunlop, 2010; Waddington et al., 2015) (Fig. 1). Furthermore, our results suggest that this event was independent of the apparent WGDs shared by all extant horseshoe crabs (Kenny et al., 2016).

### Divergence in gene function after duplication

It is thought that typically large scale duplication events such as WGD are followed by a period of gene loss (for example, only of 12% paralogs have been retained after 100 MY in *Saccharomyces cerevisiae;*Kellis et al. (2004); Wolfe and Shields (1997)), in concert with major genomic rearrangements, and that those duplicated genes that are subsequently retained are enriched in developmental genes such as those encoding transcription factors and other proteins that often act in multiprotein complexes (Davis and Petrov, 2005; Hakes et al., 2007; Papp et al., 2003a; Putnam et al., 2008; Semon and Wolfe, 2007). Our GO term enrichment analysis partially confirms a similar trend for *P. tepidariorum*, since we find, for instance, proteins related to transcriptional regulation enriched in the group of duplicates. Indeed, it is striking that vertebrates, horseshoe crabs and arachnopulmonates have retained duplicated Hox clusters and appear to be enriched in other paralogs that encode other transcription factors, suggesting that this retention pattern after WGDs is a general trend in animals.

Our study provides evidence for possible subsequent subfunctionalization and neofunctionalization among ohnologs (Force et al., 1999; Lynch and Conery, 2000; Lynch and Force, 2000; Lynch et al., 2001; Papp et al., 2003a) most likely as a result of evolutionary changes in their regulatory sequences as has been observed in the case of other WGD events (Papp et al., 2003b). This is exemplified by the diversity in the temporal and spatial expression of the *P. tepidariorum* Hox gene paralogs during embryogenesis (e.g. Fig. 9), Divergence in the expression patterns of duplicated Hox genes has been previously reported for the genes *Dfd, Scr* and *Ubx* in spiders (Abzhanov et al., 1999; Damen et al., 1998; Schwager et al., 2007) and for the posterior Hox genes *Antp, Ubx, abdA* and *AbdB* in the scorpion *C. sculpturatus* (Sharma et al., 2014b). However, these previous studies only investigated a few Hox gene families and analysis of the spatial expression of these genes was limited to later developmental stages after the appearance of limb buds. Divergence in gene expression has also been observed previously for duplicated Wnt ligand genes in *P. tepidariorum* (Janssen et al., 2010). In addition, a recent study of the two *dachshund* paralogs provided possible evidence the neofunctionalization of a duplicated gene during the evolution of a morphological novelty in spiders (Turetzek et al., 2016).

### Gene duplication and arachnid evolution

Our findings have profound implications for the evolution of chelicerates as a whole, a group whose internal phylogeny has proven extremely difficult to resolve (Sharma et al., 2014a). Focal to understanding the evolution of terrestrialization in this group are the relationships of five arachnid orders possessing book lungs. The close relationship of four of these groups, spiders, amblypygids, thelyphonids and schizomids, is generally not contested and both morphological and molecular trees place them together in a monophyletic clade, the Tetrapulmonata. The position of scorpions in the chelicerate tree is, however, much more controversial. It has been argued that their terrestrial adaptations, including the book lungs, evolved convergently to those of tetrapulmonates, whereas recent phylogenomic analyses have placed scorpions (possibly sister to Pseudoscorpiones) as the sister group to Tetrapulmonata (Regier et al., 2010; Sharma et al., 2014a). The shared paleopolyploidization event between spiders and scorpions provides further evidence that these two groups are more closely related to each other than they are to other apulmonate and non-duplicated arachnids (e.g. mites and ticks), which is in agreement with recent molecular phylogenies. This would imply a single origin of the arachnid book lungs as has been suggested previously based on detailed ultrastructural morphological analyses (Scholtz and Kamenz, 2006), raising the possibility that the ancient WGD identified here can be tested using new comparative genomic data, sampling such lineages as amblypygids, thelyphonids and schizomids.

The age of the duplication event identified here must predate the most recent common ancestor of spiders and scorpions. Molecular clock approaches vary widely on the age of arachnids, and have suggested that arachnids diversified in the Ordovician (Lozano-Fernandez et al., 2016; Rota-Stabelli et al., 2013) or in the Silurian (Sharma and Wheeler, 2014), with large confidence intervals on node age estimates that often span entire geological periods. However, the earliest stem-group spiders (the extinct order Uraraneida) date to the mid-Devonian (386 MYA; Selden et al. (2008)), whereas discoveries of Paleozoic scorpions have extended the stratigraphic range of scorpions into the Silurian (430 MYA; Waddington et al. (2015)). The arachnid fossil record thus suggests the mid-Silurian is a conservative floor age of the duplication event. A Paleozoic age of the duplication event at the base of Arachnopulmonata would make this event approximately contemporaneous with the two-fold WGD in the ancestral vertebrate (Dehal and Boore, 2005).

This reconstruction is consistent with the observation that few genes retain the ancient signal of shared duplication in both arachnopulmonates and vertebrates, and those that do often tend to be developmental patterning genes. For example, when compared to the *Drosophila melanogaster* genome, less than 5% of homologous vertebrate genes retain the 1:4 ortholog ratio expected from the vertebrate two-fold WGD event (Dehal and Boore, 2005). But included among this minority are vertebrate orthologs of Hox genes, whose duplicates have been retained and deployed for various aspects of embryonic patterning. Thus, the patterns observed in arachnopulmonate arachnids are broadly consistent with counterparts in vertebrates.

Currently, it is not possible to address the question of whether the arachnopulmonate WGD facilitated the evolution of a terrestrial life-style and the development of book lungs. Taking advantage of the annotated spider genome sequences and the practical merits of the spider model, however, future functional studies in spiders could analyze paralog sub- and neo-functionalization and gene regulatory network rewiring after duplication to clarify these questions.

### Implications for future studies on genome evolution

Much has been speculated about the long term evolutionary consequences of genome duplications, including long-standing discussions on the evolution and origin of our own lineage, the vertebrates, and the complex body plan and diverse ecological adaptations that are hallmarks of this animal group (Freeling et al., 2015; Glasauer and Neuhauss, 2014; Holland, 2013; Ohno, 1970; Semon and Wolfe, 2007). However, it has been argued that there does not appear to be an association between genome duplication and teleost diversification (Clarke et al., 2016). Furthermore, other groups that experience WGD, such as horseshoe crabs and bdelloid rotifers, did not exhibit any apparent diversification or increase in complexity following WGD, which suggests that a putative link between WGD and increased diversification, as suggested in vertebrates, may not be generalizable to other taxa (Flot et al., 2013; Nossa et al., 2014; Ricci, 1987; Rudkin and Young, 2009).

To help address the contribution of WGD to animal diversification, analyzing the outcomes of those independent “experiments” that have naturally occurred during evolutionary time is of paramount importance. Recurrent and independent cases of paleopolyploidization should be studied systematically to reveal commonalities of evolutionary forces experienced across disparate lineages. Our discovery of an ancient genome duplication event preceding the origin of spiders and scorpions helps to fill a crucial gap in the comparative studies of WGDs. Previously reported cases of paleopolyploid lineages in different eukaryotes, including both unicellular and multicellular taxa, only allowed an extremely reduced set of core orthologous genes to be compared across lineages. However, the biology of vertebrates and arachnopulmonates is in many respects very similar, sharing the gene toolkit common to most animal species, highly conserved developmental pathways and even the general layout of the basic bilaterian body plan. Thus, our results will open new research avenues, allowing the formulation of specific hypotheses about the impact of WGDs on developmental gene regulatory networks and morphological diversity by making direct comparisons and extrapolations with the vertebrate case.

## Materials and Methods

### Extraction of genomic DNA

Genomic DNA was extracted from four adult females and eight adult males of a genetically homogenous *P. tepidariorum* strain that was inbred for 15 generations and originally collected in Göttingen. All 12 animals were separated from the general stock before their final molt (to ensure that all specimens were virgin and did not contain genetic material from mating partners or developing embryos), and were starved for two weeks prior to DNA extraction (to minimize contamination from gut contents). Directly before DNA extraction, all animals were microscopically inspected to ensure they were free of external parasites (e.g. mites) and were macerated and digested in 80 mM EDTA (pH=8.0), 100 mM Tris-HCl (pH=8.0), 0.5% SDS, and 100 pg/ml proteinase K at 60°C for 2 hours. Genomic DNA was isolated from this solution by salt-chloroform extraction, precipitated with ammonium acetate and ethanol, and dissolved in water. RNA contamination was removed with RNaseA. Purified genomic DNA was precipitated with sodium acetate, washed with ethanol and dissolved in TE-buffer (10 mM Tris-HCl (pH=7.4), 1 mM EDTA) (pH=8.0)).

For the bark scorpion *C. sculpturatus*, genomic DNA was extracted from four legs, a pedipalp patella and femur, and the fourth metasomal segment of an adult wild-caught female specimen (Tucson, Arizona, USA). Extraction was performed using the Animal Blood and Tissue protocol for a Qiagen DNeasy kit, with the addition of 16 ul of RNase A (25 mg/ml). Whole body RNA was extracted from the same adult female, an adult male, and a juvenile using one leg, the telson, the fifth metasomal segment, 1/3 of the abdomen (to avoid gut contamination), and 1/2 of the cephalothorax, and a pedipalp patella. Total RNA was extracted using Trizol with the addition of glycogen.

### Genome sequencing and assembly

The house spider and bark scorpion are two of 30 arthropod species sequenced as part of the pilot project for the i5K 5000 arthropod genomes project at the Baylor College of Medicine Human Genome Sequencing Center. For all of these species an enhanced Illumina-ALLPATHS-LG sequencing and assembly strategy enabled multiple species to be approached in parallel at reduced costs. For the house spider, we sequenced five libraries of nominal insert sizes 180 bp, 500 bp, 2 kb, 3 kb and 8 kb at genome coverages of 39.2×, 35.1×, 19.7×, 49.3×, 19.3× respectively (assuming a 1.5 Gb genome size (Posnien et al., 2014)). These raw sequences have been deposited in the NCBI SRA: BioSample ID SAMN01932302. For the bark scorpion, we sequenced four libraries of nominal insert sizes 180 bp, 500 bp, 3 kb and 8 kb at genome coverages of 102.1×, 25.6×, 35.2×, 39.0× respectively (assuming a 900 Mb genome size). These raw sequences have been deposited in the NCBI SRA: BioSample SAMN02617800.

To prepare the 180 bp and 500 bp libraries, we used a gel-cut paired-end library protocol. Briefly, 1 μg of the DNA was sheared using a Covaris S-2 system (Covaris, Inc. Woburn, MA) using the 180 bp or 500 bp program. Sheared DNA fragments were purified with Agencourt AMPure XP beads, end-repaired, dA-tailed, and ligated to Illumina universal adapters. After adapter ligation, DNA fragments were further size-selected on an agarose gel and PCR-amplified for 6 to 8 cycles using the Illumina P1 and Index primer pair and Phusion^®^ High-Fidelity PCR Master Mix (New England Biolabs). The final library was purified using Agencourt AMPure XP beads and quality assessed by Agilent Bioanalyzer 2100 (DNA 7500 kit) to determine library quantity and fragment size distribution before sequencing.

Long mate pair libraries with 2 kb, 3 kb and 8 kb insert sizes were constructed according to the manufacturer’s protocol (Mate Pair Library v2 Sample Preparation Guide art # 15001464 Rev. A PILOT RELEASE). Briefly, 5 pg (for 2 and 3-kb gap size library) or 10 μg (8-10 kb gap size library) of genomic DNA was sheared to desired size fragments by Hydroshear (Digilab, Marlborough, MA), then end repaired and biotinylated. Fragment sizes between 1.8 - 2.5 kb (2 Kb) 3 - 3.7 kb (3 kb) or 8-10 kb (8 kb) were purified from 1% low melting agarose gel and then circularized by blunt-end ligation. These size selected circular DNA fragments were then sheared to 400 bp (Covaris S-2), purified using Dynabeads M-280 Streptavidin Magnetic Beads, end-repaired, dA-tailed, and ligated to Illumina PE sequencing adapters. DNA fragments with adapter molecules on both ends were amplified for 12 to 15 cycles with Illumina P1 and Index primers. Amplified DNA fragments were purified with Agencourt AMPure XP beads. Quantification and size distribution of the final library was determined before sequencing as described above.

Sequencing was performed using Illumina HiSeq2000 generating 100 bp paired end reads. Reads were assembled using ALLPATHS-LG (v35218) (Gnerre et al., 2011) and further scaffolded and gap-filled using Atlas-Link (v.1.0) and Atlas gap-fill (v.2.2) (https://www.hgsc.bcm.edu/software/). For *P. tepidariorum*, this yielded an assembly size of 1443.9 Mb with 263,833 contigs with an N50 of 10.1 kb and, after scaffolding and gap closing, 31,445 scaffolds with an N50 of 465.5 kb. Approximately 2,416 million reads (96.9x sequence coverage) are represented in this assembly of the *P. tepidariorum* genome. The assembly has been deposited in the NCBI: BioProject PRJNA167405 (Accession: AOMJ00000000).

For the *C. sculpturatus* this yielded an assembly size of 926.4 Mb with 214,941 contigs with an N50 of 5.1 kb and, after scaffolding and gap closing, 10,457 scaffolds with an N50 of 342.5 kb. The final assembly has been deposited in the NCBI: BioProject PRJNA168116.

### Dovetail assembly

#### Chicago library preparation

To improve the *P. tepidariorum* assembly further we used *in vitro* contact genomics (Flot et al., 2015) based on the Chicago method (Dovetail Genomics, Santa Cruz, CA; (Putnam *et al.* 2016)). A Chicago library was prepared as described previously (Putnam et al, 2016). Briefly, ≥ 0.5 μg of high molecular weight genomic DNA ≥50 kb mean fragment size was extracted from a female *P. tepidariorum*, reconstituted into chromatin *in vitro*, and fixed with formaldehyde. Fixed chromatin was then digested with *MboI* or DpnII, the 5’ overhangs were filled in with biotinylated nucleotides, and then free blunt ends were ligated. After ligation, crosslinks were reversed and the DNA purified from protein. Purified DNA was treated to remove biotin that was not internal to ligated fragments. The DNA was sheared to ~350 bp mean fragment size, and sequencing libraries were generated using NEBNext Ultra enzymes and Illumina-compatible adapters. Biotin-containing fragments were then isolated using streptavidin beads before PCR enrichment of the library.

#### Scaffolding the draft genome with HiRise

The *P. tepidariorum* draft genome in FASTA format (1443.9 Mb with a scaffold N50 of 465.5 kb), the shotgun sequences (from approximately 2,416 million Illumina reads (see above)) and the Chicago library sequence (187 million read pairs from Illumina HiSeq 2500 2X100bp rapid run) in FASTQ format were used as input data for HiRise, a software pipeline designed specifically for using Chicago library sequence data to assemble genomes (Putnam *et al.* 2016). Shotgun and Chicago library sequences were aligned to the draft input assembly using a modified SNAP read mapper (http://snap.cs.berkeley.edu). The separations of Chicago read pairs mapped within draft scaffolds were analyzed by HiRise to produce a likelihood model, and the resulting likelihood model was used to identify putative misjoins and score prospective joins. After scaffolding, shotgun sequences were used to close gaps between contigs. This resulted in 16,542 superscaffolds with an N50 of 4,050 kb.

### Genome annotation

#### P. tepidariorum

The *P. tepidariorum* genome assembly (pre-Dovetail) was annotated using version 2.7 of AUGUSTUS (Stanke et al., 2008). AUGUSTUS constructed genes from evidence such as the RNA-Seq alignments - here called hints - but also uses statistical models for *ab initio* prediction. The parameters for the statistical models of *P. tepidariorum* genes were estimated on a training set of gene structures. Several steps of parameter estimation, prediction, visual quality control on a genome browser and parameter tuning were performed.

*P. tepidariorum* transcript alignments were generated using available RNA-Seq libraries (Posnien et al., 2014): 1,040,005 reads from 454-sequencing of *P. tepidariorum* embryonic stages, two RNA-Seq libraries from Illumina-sequencing of embryonic stages (333,435,949 and 602,430 reads), and two RNA-Seq libraries from Illumina-sequencing of post-embryonic stages (294,120,194 read and 317,853 reads). In addition, we downloaded all *P. tepidariorum* ESTs (Oda et al., 2007) and protein sequences available in Genbank. The assembly was repeat-masked using RepeatMasker (version 1.295, (Smit et al., 1996-2010)) and TandemRepeatFinder (version 4.07b, (Benson, 1999)) based on a de novo repeat library compiled with RepeatScout (version 1.0.5, (Price et al., 2005)). 46% of the bases were masked as repeats.

*P. tepidariorum-specific* parameters of AUGUSTUS were estimated iteratively. An initial training set of genes was generated with PASA (release 2012-06-25, (Haas et al., 2008)) using the ESTs only. This yielded 851 genes that were used to estimate the first set of parameters of AUGUSTUS for the coding regions of genes. Additionally, eukaryotic core proteins were predicted in the masked assembly with CEGMA (version 2.4.010312) (Parra et al., 2007) and yielded 103 hints for CDS to AUGUSTUS which were then used in the training stage predictions. With these initial parameters and integrating the evidence from transcriptome data, AUGUSTUS was used to annotate the masked assembly genome-wide. We then extracted another training gene set from the genome-wide prediction according to the following procedure: RNA-Seq reads from 454 and Illumina sequencing were mapped against predicted transcripts using GSNAP (version 2013-06-27) (Wu and Nacu, 2010). (i) Only genes with 100% RNA-Seq alignment coverage were taken. (ii) We mapped the proteins from the database UniRef50 (version UniProt Release 2013 06) (2014) against predicted proteins using BLASTP (version 2.2.25) (Altschul et al., 1990) and kept only fully covered transcripts. The genes in the intersection of both sets - that is genes fulfilling constraints (i) and (ii) simultaneously - were used for a second iteration of parameter training. The UTR parameters of AUGUSTUS were only trained once when other parameters had already become stable.

RNA-Seq reads from 454 and Illumina sequencing were mapped against the masked assembly using GSNAP (version 2013-06-27) (Wu and Nacu, 2010). The evidence from transcriptome data, protein homology and repeats was input to AUGUSTUS as a ‘hints’ file. The spliced alignments of the RNA-Seq reads using GSNAP resulted in 272,816 unique intron hints and further hints on exonic parts from transcribed regions. Further, we obtained 97,785 hints from ESTs (not only for CDS) using BLAT (version v. 35×1) (Kent, 2002). The roughly 2.1 million repeat-masked regions were used as ‘nonexonpart’ hints in the annotation, moderately penalizing the prediction of exons overlapping repeats. Consecutive gene sets were computed utilizing AUGUSTUS to stepwise improve prediction accuracy and reliability of the final gene set release referred to as aug3. All extrinsic hint data were incorporated into this last prediction. Allowing the occurrence of alternative transcripts in the results, the final gene set aug3 was then generated using the call:

Augustus-species=parasteatoda -alternatives-from-evidence=true … --UTR=on --hintsfile=all.hints --extrinsicCfgFile=extrinsic.P.E.RM.cfg genome_masked.fa

The RNA-Seq data coverage was quantified using the transcript quantification tool express (Roberts and Pachter, 2013), which estimates FPKM-values (FPKM: fragments per kb of transcript per million mapped reads at transcript level), thereby quantifying the pooled abundances of the predicted transcripts in the RNA-Seq data.

The aug3 gene models were transferred to the Dovetail genome assembly using Exonerate v2.2 (Slater and Birney, 2005) with the command --model protein2genome --bestn 1 -- showtargetgff YES. The resulting GFF files were converted into protein sets from the corresponding Dovetail genome fasta file.S

The Trinotate annotation pipeline (Release 2.0.2; https://trinotate.github.io/) was used for the functional annotation of the aug3 protein predictions following the standard procedure. In summary, the predicted peptide sequences of the aug3 annotation were blasted against UniRef90 and SwissProt databases with E <= 0.05 and keeping only the best hit. HMMER (version 3.1b1; http://hmmer.org/) was used to search the Pfam database to predict protein domains. All Blast searches were run in parallel on a high performance computer cluster utilizing the perl script HPC GridRunner (v1.0.2; http://hpcgridrunner.github.io/). The Blast and protein domain predictions were stored in a predefined sqlite (version 3.8.8.3; https://www.sqlite.org/) database. Trinotate was used to export a final report that contains the best Blast hits, protein domain predictions and GO categories extracted from the Blast result and the Pfam domain prediction for each of the aug3 predictions (Table S16).

The final annotated gene set contains 27,990 genes and 31,186 transcripts. 85% of the predicted *P. tepidariorum* proteins had homology support derived from a BLASTP search against the UniRef50 data (E value <= 10^−5^). Transcript quantification from the RNA-Seq data (using estimates of FPKM-values: fragments per kb of transcript per million mapped reads at transcript level (Roberts and Pachter, 2013)) showed that 29,966 (93%) of predicted transcripts had transcriptome support at FPKM ≥ 0.034 and 26,381 (82%) of predicted transcripts had transcriptome support at FPKM ≥ 0.34. In the final gene set only 1.1% of the predicted transcripts had neither homology nor transcriptome support at an FPKM threshold of less than 0.034. The annotated *P. tepidariorum* genome is available at https://i5k.nal.usda.gov/JBrowse-partep.

#### C. sculpturatus

The *C. sculpturatus* genome was annotated using MAKER (Campbell et al., 2014) with RNA-Seq reads generated from a juvenile (http://www.ncbi.nlm.nih.gov/sra/SRX911075), adult female (http://www.ncbi.nlm.nih.gov/sra/SRX911064), and adult males

(http://www.ncbi.nlm.nih.gov/sra/SRX911079).

### Analysis of duplicated genes

#### Classification of duplicates using MCScanX

The data used to perform these analyses were, for *P. tepidariorum*, the aug3 version; and for *C. sculpturatus*, the 0.5.53 version of the MAKER annotation available at ftp://ftp.hgsc.bcm.edu/I5K-pilot/Barks_corpion/. The same analysis was also performed on the *Limulus polyphemus* genome (ftp://ftp.ncbi.nih.gov/genomes/Limulus_polyphemus/) as a comparison.

Out of the 32,949 gene models in the aug3 annotation of the *P. tepidariorum* genome (resulting from the transfer of the aug3 annotation on the Dovetail scaffolds), only the main transcript of each gene was retained, yielding a set of 28,746 gene models. This list was further shortened by removing all instances of 755 gene models that had become artefactually duplicated during the annotation transfer process from aug2 to aug3, resulting in a final set of 27,203 gene models. All of the 30,465 gene models in the *C. sculpturatus* annotation were retained for the synteny analyses. Finally, out of the 23,287 annotated proteins of *L. polyphemus*, 21,170 were retained for the synteny analyses after filtering out annotated isoforms of the same genes (based on their identical start and end positions).

Hits within and between gene sets were catalogued using BLASTP using an e-value threshold of 10^−10^ and keeping only the 5 best hits as recommended in the instruction manual of MCScanX (Wang et al., 2012). Then MCScanX was used with default parameters to classify genes into four categories: singletons (i.e., genes without any duplicate); dispersed (duplicates occurring on different scaffolds); proximal (duplicates occurring on the same scaffold less then 10 genes apart); distal (duplicates occurring more than 10 genes apart on a same scaffold); and segmental (block of at least 5 collinear genes separated by less than 25 gene missing on one of the duplicated regions).

#### Orthology assessment of arthropod genomes

To investigate the extent of gene duplication in *P. tepidariorum* and *C. sculpturatus*, we compared these two genomes to those of other four arthropods with no demonstrable evidence of a WGD. These non-arachnopulmonate taxa were another chelicerate (the tick *I. scapularis*) and three mandibulates (the flour beetle *Tribolium*, the crustacean *Daphnia pulex*, and the centipede *Strigamia maritima*). Predicted peptides sets (aug3) were used as inputs, and redundancy reduction was done with CD-HIT (Fu et al., 2012) to remove the variation in the coding regions of genomes attributed to allelic diversity R (>99% sequence similarity). Peptide sequences with all final candidate ORFs were retained as fasta files. We assigned predicted ORFs into orthologous groups across all samples using OMA stand-alone v.0.99u (Altenhoff et al., 2013; Altenhoff et al., 2011) discarding sequences less than 50 sites in length. All-by-all local alignments were parallelized across 400 CPUs. Orthology mapping of spider and scorpion genes that could be mapped to a mandibulate or tick counterpart was conducted using custom Python scripts on the OMA output.

To assess the possibility of incorrect orthology assessment stemming from algorithmic error, we identified the intersection of the OMA output (based on whole genomes) and a set of orthologs found to occur in single-copy across Arthropoda, as benchmarked in the BUSCO-Ar database of OrthoDB (Waterhouse et al., 2013). The BUSCO-Ar set of the flour beetle *T. castaneum* was selected as the reference genome for the BUSCO set.

In a separate and subsequent analysis, three additional taxa (genomes of the horseshoe crabs *L. polyphemus, Tachypleus gigas*, and *Carcinoscorpius rotundicauda*) were added to the taxa in the principal OMA run, with all other procedures as specified above.

#### Analysis of gene tree topologies from six-genome dataset

From the output of the OMA analysis of six arthropod genomes, we extracted a subset of orthogroups wherein exactly two spider paralogs were present for one *T. castaneum* ortholog (i.e., 1:2 orthology). *T. castaneum* was chosen as the reference genome in comparative analyses both for the quality of its assembly and for its archetypal gene content among Arthropoda. Gene trees for this subset of orthogroups were inferred to examine the topological relationship between homologous sequences of arachnopulmonate and non-arachnopulmonate taxa. These orthogroups were aligned using MUSCLE v.3.8 (Edgar, 2004) and ambiguously aligned regions were culled using GBlocks v.0.91b (Castresana, 2000) using the commands -b3=8 (maximum of eight contiguous non-conserved positions), -b4=10 (minimum of ten positions in a block), and -b5=h (gap positions allowed for a maximum of half the sequences). Maximum likelihood analyses were conducted using the LG + Γ model with four rate categories (Le et al., 2012; Yang, 1996) and 500 independent starts in RAxML v. 7.3.0 (Stamatakis, 2006).

We characterized whether the resulting tree topologies corresponded to Hypothesis 1 (common duplication in the most recent common ancestor (MRCA) of spiders and scorpions), Hypothesis 2 (lineage specific duplication events in each of spiders and scorpions), an indeterminate tree topology (corresponding to neither scenario, typically due to the non-monophyly of the outgroup taxa), or an uninformative tree topology (due to the lack of any scorpion paralogs). Cases where the two spider paralogs formed a grade with respect to a single scorpion paralog were additionally classified as partially congruent with Hypothesis 1. The set of gene trees either partially or fully congruent with Hypothesis 1 is henceforth termed “Tree Set 1”. Alignments and gene tree files are available on request.

#### Analysis of gene tree from nine-genome dataset

To infer the relationship between arachnopulmonate and xiphosuran paralogs, from the OMA analysis of nine genomes (the six genomes above; *L. polyphemus*; *T. gigas; C. rotundicauda*) we separately extracted another subset of orthogroups wherein two, three, or four horseshoe crab paralogs from any of the three horseshoe crab genomes were detected for one *T. castaneum* ortholog (i.e., 1:2, 1:3, or 1:4 orthology). We inferred gene trees with the approach specified above. We again distinguished two scenarios: (a) separate WGD events in the MRCA of Arachnopulmonata and Xiphosura (Hypothesis 3), and (b) a common WGD event in the MRCA of all Chelicerata (Hypothesis 4). Cases where ancient paralogy was detected in Xiphosura alone (and not Arachnopulmonata) were classified as partially congruent with Hypothesis 3. The set of gene trees either partially or fully consistent with Hypothesis 3 was termed “Tree Set 2”. Alignments and gene tree files are available on request.

#### *Identification of paralog pairs in* P. tepidariorum *and other chelicerates*

Putative families of homologous protein-coding genes were identified for 31 chelicerate species and a myriapod (Table S7). Protein sequences from the publically available translated coding sequences were also used. Otherwise, transcripts were translated with Transdecoder (Haas et al., 2008). For translated sequences with >95% identity, only the single longest protein was retained for further analyses. For transcripts assembled by Trinity (Grabherr et al., 2011), the longest transcript per “contig” was retained (Trinity often generates multiple transcripts associated with a single “contig”, thought to represent isoforms).

We grouped genes into families using a modified version of the method applied in the Phytozome project described in (Goodstein et al., 2012), with either *P. tepidariorum* or *C. sculpturatus* translated genes used as a seed. In short, homologous protein pairs were identified using all-versus-all BLASTP comparisons of the 32 arthropod species with an e-value < 1e-3 cutoff (Altschul et al., 1990). A global alignment score was calculated for each homologous pair using the Needleman-Wunsch algorithm with the Blosum62 matrix. We then used the Needleman-Wunsch score between *P. tepidariorum* (or *C. sculpturatus*) protein sequences and the rest of the sequences to seed the gene families in a three-step process. First, for each non-*P. tepidariorum* protein, the *P. tepidariorum* protein with the highest Needleman-Wunch score was identified. Second, all the *non-Parasteatoda* proteins with the same best-scoring *P. tepidariorum* protein were grouped with the *P. tepidariorum* protein. Third, all the groups were combined that contained *P. tepidariorum* proteins determined to be homologous to each other based on a BLASTP alignment with an e-value < 1e-3. The same three-step process was repeated to identify *C. sculpturatus* - seeded gene families.

For each gene family, the protein sequences were multiply aligned using MUSCLE (Edgar, 2004). The multiple alignments were trimmed by removing all the bounding alignment positions that added more gaps than sequence by a custom Perl script. Entire protein sequences were removed from the alignment if the sequence had gaps in more than 25% of the aligned positions. For the *P. tepidariorum*-seeded gene families, only those containing at least one *P. tepidariorum* protein and four additional sequences were retained for further analyses. Within the retained families, poorly aligned columns were removed using TrimAL under a “strict-plus” setting, which optimizes the signal to noise ratio in the multiple alignment (Capella-Gutierrez et al., 2009). The protein alignments were then used to guide nucleotide alignments by replacing the amino acids with their encoding transcript sequences.

Protein alignments were used to infer gene trees with TreeBeSt (http://treesoft.sourceforge.net/treebest.shtml). TreeBeST searches for an optimal gene tree given a species tree (we used the phylogeny in Fig. S6) and identifies duplication and speciation events within the optimal tree. Branch lengths were calculated for the optimal TreeBeSt tree using maximum likelihood (PhyML type search) with the HKY model of evolution (Hasegawa et al., 1985). Alignments and gene tree inferences were repeated for the *C. sculpturatus*-seeded gene families.

#### Molecular distance of duplication and speciation events

We estimated the molecular distance of a *P. tepidariorum* (or *C. sculpturatus*) duplication or speciation node in *P. tepidariorum* (or *C. sculpturatus*)-seeded families by averaging the branch lengths in TreeBeST trees from the node to all its *P. tepidariorum* (or *C. sculpturatu*s) descendants. We similarly estimated molecular distance of other species’ duplication nodes by averaging the branch length from the node to all of the species of interest’s descendants. Distributions of molecular distances were estimated and statistical tests for goodness of fit calculated in R.

#### *Ascertaining GO Term enrichment in* P. tepidariorum *paralog pairs*

GO Terms were imputed to the *P. tepidariorum* AUGUSTUS gene models (aug3) through comparisons to the UniRef50 protein set by BLASTP comparisons using a cut-off of 1e-5. The GO Terms of its closest UniRef by e-value with documented GO Terms were assigned to a gene model via a custom perl script, with GO Slim values derived using GOSlimViewer (available at http://agbase.msstate.edu/cgi-bin/tools/goslimviewer_select.pl (McCarthy et al., 2006). Enrichment of GO Terms within gene families was ascertained using Fisher’s Exact Test.

#### Synteny analyses

A genome-scale synteny analysis of the *P. tepidariorum* scaffolds was conducted using the program SatsumaSynteny (Grabherr et al., 2010). This approach does not rely on the annotation and can detect weak, degraded signals of synteny such as signatures of ancient WGDs that were followed by numerous rearrangements. For visualization, we selected only the 100 scaffolds for which the number of hits detected by Satsuma was maximal; in a second time, this list was further reduced to the set of 39 scaffolds that exhibited the greatest number of hits with each other. An Oxford grid (Edwards, 1991) was drawn using the tool orthodotter (available online from http://www.genoscope.cns.fr/externe/jmaurv/orthodotter.tar.gz). and a circular plot was drawn using Circos (Krzywinski et al., 2009).

For the synteny analysis of selected developmental genes, their nucleotide sequences were first downloaded from NCBI (Accession numbers are given in Table S5). BLASTN searches against the Augustus 3 gene set were used to identify the best aug3 prediction and BLASTN searches against the Dovetail assembly (Assembly 2.0) were used to identify their respective scaffold.

All 148 precursor microRNAs sequences for *P. tepidariorum* (Leite et al., 2016), with the inclusion of flanking sequences 20 bp up and down stream, were BLASTN-searched in the Dovetail assembly to identify scaffold ID and position from the best matches. The scaffolds and positions of *C. sculpturatus* microRNAs from Leite et al. (2016) were used.

#### Homeodomain and Hox gene annotation

To identify possible homeobox genes in *P. tepidariorum* and *C. sculpturatus*, the complete set of homeobox sequences from HomeoDB (Zhong et al., 2008; Zhong and Holland, 2011), those identified previously in the scorpion *Mesobuthus martensii* (Di et al., 2015) and the *P. tepidariorum* gene *prospero* (Accession: BAE87100.1) were BLASTP (version 2.4.0+) (Camacho et al., 2009) searched against the *P. tepidariorum* AUGUSTUS (aug3) and *C. sculpturatus* MAKER protein predictions. All blast hits were scanned for the presence of homeobox domains and other functional domains with the CDD search tool (Marchler-Bauer et al., 2015). Hits that contained at least one homeobox sequence were manually checked for the completeness of the homeobox domain sequence. The homeobox genes were annotated and classified based on Holland et al. (2007).

To identify the location of Hox genes on genomic scaffolds of *P. tepidariorum, Latrodectus hesperus, S. mimosarum, A. geniculata, C. sculpturatus, I. scapularis* and genomic contigs of *M. martensii* were searched for Hox genes with tblastx BLAST (version 2.2.28+) (Camacho et al., 2009) using published chelicerate Hox gene sequences. Scaffolds or contigs containing blast hits to Hox genes were extracted and intron-exon boundaries were hand-annotated in Geneious (version 7) (Kearse et al., 2012) with the help of sequenced transcriptomes, sequences obtained by RACE PCR experiments (in the case of *P. tepidariorum)*, cloned Hox gene sequences (in the case of *C. sculpturatus*), or by comparison between the chelicerate sequences. In case of additional splice variants containing additional small exons, the shortest version consisting only of two exons was used for the analysis. Naming of Hox genes followed orthologies to already published Hox genes in *C. salei* and *P. tepidariorum* for the spider sequences or, in the case of the scorpions, orthologies to published *C. sculpturatus* sequences.

#### Hox gene alignments and topological tests of gene trees

Nine Hox class genes were used as test cases for distinguishing two scenarios: (a) common duplication in the MRCA of spiders and scorpions (Hypothesis 1) and (b) lineage specific duplication events in each of spiders and scorpions (Hypothesis 2). The single remaining Hox class gene (*fushi tarazu*) did not possess the minimum requirement—inclusion of two paralogs each of a spider and a scorpion species—and thus was not dispositive in topological tests. Peptide sequence alignments were constructed using MUSCLE v. 3.8 (Edgar, 2004) and alignment ends manually trimmed, such that either terminus of the alignment sampled at least half of each alignment’s terminals. Preliminary efforts using outgroup taxa have demonstrated little statistical power resulting from rooting trees, due to large phylogenetic distances between arachnopulmonates and arachnid outgroups (e.g., harvestmen, pycnogonids; (Sharma et al., 2014b)), as well as accelerated evolution in other potential outgroup taxa (e.g., mites, ticks; (Sharma et al., 2014a)). For this reason, outgroup-free tests were conducted using spider and scorpion sequences only.

Maximum likelihood analyses were conducted using the LG + Γ model with four rate categories (Le et al., 2012; Yang, 1996) and 500 independent starts in RAxML v. 7.3.0 (Stamatakis, 2006). To compare tree likelihoods of unconstrained runs to Hypothesis 2, a constraint tree was imposed for each Hox class enforcing mutual monophyly of spider and scorpion sequences, and the best tree topology was selected from 500 independent starts under the scenario of lineage-specific duplications.

## Embryos, *in situ* hybridization and imaging

*P. tepidariorum* embryos were obtained from laboratory cultures in Oxford, UK, Cambridge, MA, USA and Cologne, Germany. RNA was extracted from embryos of stages 1-14 using either Trizol (Life Technologies) or Qiazol (Qiagen) and cDNA was synthesized with SuperscriptIII (Life Technologies). Probe templates were either synthesized by PCR using TOPO pCR4 vectors containing cloned RACE fragments of Hox genes (RACE was performed with the Marathon RACE kit or SMART RACE cDNA kit (Clontech)), or they were generated by adding T7 binding sites to RT-PCR fragments as described by (Sharma et al., 2012). Primer sequences used for the RT-PCR fragments were based on the *P. tepidariorum* transcriptome (Posnien et al., 2014) and genome sequences. The origin of gene fragments and primers is available on request. Embryos were fixed and probe synthesis and in situ hybridizations were carried out as described previously (Akiyama-Oda and Oda, 2003; Prpic et al., 2008). The anti-DIG antibody was pre-absorbed over night at 4°C with mixed-stage embryos. Stained embryos were staged according to Mittmann and Wolff (2012) and imaged using a Leica stereoscope fitted with a Zeiss AxioCam MRc. Images were processed in Photoshop CS4 or CS6.

## Acknowledgements

This work was supported by NIH grant NHGRI U54 HG003273 to RAG, the National Science Foundation (IOS-0951886 to NAA), a Leverhulme visiting fellowship for EES (VF-2012-016), funding and PhD studentships (DJL, LG and AS) from Oxford Brookes University, and a Conselho Nacional de Desenvolvimento Científico e Tecnológico (CNPq) scholarship to CLBP. N-MP was funded by the Deutsche Forschungsgemeinschaft (grant numbers PR 1109/4-1, PR 1109/7-1 and

PR 1109/6-1 to N-MP). Additional financial backing has been received from the Göttingen Graduate School for Neurosciences, Biophysics and Molecular Biosciences (GGNB), the Göttingen Center for Molecular Biosciences (GZMB), and the University of Göttingen (GAU). NT is supported by a Christiane-Nüsslein-Volhard-Foundation fellowship and a "Women in Science" Award by L’Oréal Deutschland and the Deutsche UNESCO-Kommission. NP has been funded by the Volkswagen Foundation (project number: 85 983) and the Emmy Noether Programme of the Deutsche Forschungsgemeinschaft (grant number: PO 1648/3-1).

## Author contributions

APM, NP, N-MP, HO, WGMD, JC, and SR conceived the project. Library construction, genome sequencing and assembly: N-MP SR, SD, SLL, HC, HVD, HD, YH, JQ, SCM, DSTH, KCW, DMM, RAG, VLG and JC. Gene duplication analyses: TC, PPS, NAA, JFF. Hox gene analysis: EES, PPS, DJL, CS. *Parasteatoda* genome data was analyzed by all authors. The manuscript was written by APM, EES, PPS, TC, NAA, MS, TW, NP, N-MP JFF and SR with the help of all other authors. We thank Wim Damen for all his support and mentoring, and for many productive discussions about spider genetics and development.

---

**Figure S1. Duplicated genes: 3 gene blocks**

**Figure S2. Duplicated genes: 4 gene blocks**

**Figure S3. Duplicated genes: 5 gene blocks**

**Figure S4. Gene tree analysis of individual Hox genes**

Blue branches indicate scorpion sequences; red branches indicate spider sequences. Gene trees are shown for *labial* **(A)**, *proboscipedia* **(B)**, *Deformed* **(C)**, *Sex combs reduced* **(D)**, *Antennapedia* **(E)**, *Ultrabithorax* **(F)**, *abdominal-A*, **(G)**, and *Abdominal-B* **(H)**. Log likelihood values are provided in Figure 4. Due to insufficient sequences for hypothesis testing, gene trees were not inferred for *Hox3-1* and *fushi tarazu.* Abbreviations: *Pt Parasteatoda tepidariorum, Lh Latrodectus hesperus, Sm Stegodyphus mimosarum, Ag Acanthoscurria geniculata, Cs Centruroides sculpturatus, Mm Mesobuthus martensii.*

**Figure S5. Synteny of 39 *P. tepidariorum* scaffolds with greatest number of reciprocal hits**

Circos plot of the subset of 39 scaffolds presenting the greatest numbers of hits on one another (as detected using SatsumaSynteny). One unit on the perimeter represents one Mbp.

**Figure S6. Phylogeny of 31 arthropod species used in comparison with *P. tepidariorum***

Species relationships are based on Bond et al. (2014); Regier et al. (2010); Sharma and Giribet (2014). The number of genes within *P. tepidariorum-seeded* gene families are shown parenthetically (n=), the number of speciation nodes observed in the gene families between *P. tepidariorum* and the other species (N=), and the median HKY distance between each speciation node and *P. tepidariorum* (HKY=) descendants are shown at the node.

**Figure S7. HKY distance distributions and Gaussian mixture models of duplication nodes from *P. tepidariorum-seeded* gene families for eight arachnid species**

The HKY distances for duplication nodes were calculated as the mean HKY branch length from the duplication node to each of the descendent genes in the species of interest (A-H). In panel (I) the distribution is for all the duplication nodes with at least one *P. tepidariorum* descendant, using the mean HKY distance from the node to *P. tepidariorum* descendants. For each panel, the best match to one of five distributions (Uniform, exponential (G), or a Gaussian mixture model with 1, 2 (H), or 3 (A-F, I) distributions) is shown. The Gaussian mixture models were seeded with Gaussian mean and standard deviations estimated from the *P. tepidariorum* duplication nodes (Figure 6a).

**Figure S8. HKY distance distributions of all *P. tepidariorum* speciation nodes in *P. tepidariorum-seeded* gene families.**

The distributions are for mean HKY branch lengths from the *P. tepidariorum* speciation node to the descendant *P. tepidariorum* genes.

**Figure S9. Tandem duplications are abundant in young duplication events, but rare in older duplication events**

HKY distance distributions and Gaussian mixture models of duplication nodes from *P. tepidariorum-seeded* gene families mirror those in Fig 6a, but paralog pairs that are found in tandem (within five genes of each other on the same scaffold) are now shown in light grey, and dispersed paralogs in dark grey.

**Figure S10. *P. tepidariorum proboscipedia-1* expression**

*pb-1* expression appears later, in a broader domain, and at much lower levels compared to *pb-2* (Fig. S11). In contrast *to pb-2, pb-1* expression is located only in the mesoderm. Weak expression of *pb-1* emerges at stage 8 in a domain spanning from Pp - L4 (*A*). At stage 8.2 *pb-1* also visible in O1 (arrow) (*B*). At stages 9 and 10, *pb-1* is expressed in the mesoderm of the outgrowing appendages of Pp-L4, as well as in dorsolateral parts of the neuroectoderm of segments Pp-O1 (arrows) (*C* and *D*). Each panel (except A) shows the same embryo, viewed laterally (left) and ventrally (center, right). Anterior is to the left. Abbreviations: Pp pedipalpal segment, L walking leg segments, O opisthosomal segments.

**Figure S11. *P. tepidariorum proboscipedia-2* expression**

Differently from *pb-1* (Fig. S10), *pb-2* expression appears earlier, in an initially much narrower domain that is restricted to the presumptive pedipalpal segment, and is mostly ectodermal. *pb-2* is first expressed in a thin stripe in the anterior of the embryo at stage 7 (*A* and *B*). The stripe broadens and as segmentation commences, it is found in the pedipalpal segment, and additionally, four weaker stripes start to appear in L1-L4 (*C*). The expression in the pedipalpal segment remains strongest, when by the end of stage 8.1, another stripe of weak *pb-2* expression is found in O1 (arrow) (*D* and *E*). During stage 9 and 10, *pb-2* expression is found in the mesoderm of the outgrowing pedipalp and walking legs, but also ectodermally in the distal tip of the outgrowing appendages of Pp-L4 (caret) (*F-H*). In the nervous system, *pb-2* is expressed most strongly in the Pp segment, starting at stage 9 (arrowhead), but it is also present in a smaller lateral domain of L1-O1 (arrow) (*F-H*). Each panel shows the same embryo, viewed laterally (left) and ventrally (right, or center and right in *E-H*). Anterior is to the left. Abbreviations: see Figure S10

**Figure S12. *P. tepidariorum Hox3-1* expression**

*Hox3-1* is not expressed before stage 8, when expression starts in broad segmental stripes from Pp to L4, most likely in the mesoderm (*A*). Expression is strongest in L1 and L4 (*A* and *B*). In stage 9 and 10 embryos, *Hox3-1* is expressed in the mesoderm of the outgrowing limb buds, except for the pedipalps, which show a broad ring of expression (white arrow in *C*) that later refines into to rings and additional expression at the tip of the pedipalp (*D-F*). Additionally expression extends anterior-ventral to the limb buds in a triangular shape at stage 9 (arrowhead in *C*). This expression vanishes by stage 10. Instead *Hox3-1* is now also expressed in the pedipalpal and walking leg segments in segmental groups of cells in the medial neuroectoderm (arrowheads in *D-F*). In stage 11 embryos, *Hox3-1* is additionally expressed in dots in the ventral neuroectoderm of every opisthosomal segment (carets in *F*). Furthermore, a dot of expression can be found in the first opisthosomal limb bud (arrow in *F*). Note that we did not detect any expression for *Hox3-2.* Embryos in *A* and *B* are shown laterally, embryos *C-E* are shown laterally on the left and ventrally on the right. *F* shows ventral views of the head region (left) and the opisthosoma (right) of an embryo at a similar stage as the embryo in *E*. Anterior is to the left. Abbreviations: see Figure S10.

**Figure S13. *P. tepidariorum Deformed-1* expression**

*Dfd-1* is expressed much earlier and more strongly than *Dfd-2* (Fig. S14). *Dfd-1* is first expressed at stage 4, in almost all cells but the rim (arrowhead) of the germ disc (*A*). During stage 6, cells not expressing *Dfd-1* at the outer rim, the future anterior, have multiplied (arrowheads)(*B*). Later during stage 6, the expression clears from the posterior (former center of the germ disc, arrow) and the expression starts to form a broad stripe (bracket) (*C*). This uniform stripe of expression subdivides first into two stripes with lower expression between these stripes during stage 7 and then the posterior domain splits into two more stripes and in between the anterior and posterior domains a new stripe gets inserted so that there are now four stripes of *Dfd-1* expression (asterisks) (*D*). These 4 stripes are later located in the four walking leg segments at stage 8 (*E*). In stage 9-11 embryos, *Dfd-1* is strongly expressed in the outgrowing walking legs with strongest expression in the tips of the legs (white carets) (*E-G*) and in the neuroectoderm (arrowheads in *F* and *G*). Each panel (except for *A, B, C*) shows the same embryo, viewed laterally (left) and ventrally (right). Anterior is to the left in all panels. Abbreviations: See Fig. S10.

**Figure S14. *P. tepidariorum Deformed-2* expression**

In contrast to *Dfd-1* (Fig. S13), *Dfd-2* is expressed later, at much lower levels and its later expression is restricted entirely to the neuroectoderm. *Dfd-2* expression is first detected in a weak, broad stripe in the developing germ band (bracket) (*A*). This stripe can be allocated to the first two walking leg segments at stage 8 (*B*). The anterior of the segments is stained stronger than the posterior. Additional, but much weaker expression can be seen in the L3 and L4 segments. At stage 9, *Dfd-2* is predominantly expressed in the ventral neuroectoderm (*C*). The anterior and posterior expression domain is marked by arrowheads in *B* and *C* respectively. Each panel shows the same embryo viewed laterally (left) and ventrally (right). Anterior is to the left. Abbreviations: See Fig. 10.

**Figure S15. *P. tepidariorum Sex combs reduced-1* expression**

Compared to *Scr-2* (Fig. S16), *Scr-1* is expressed one stage earlier and initially in a broader domain. Its anterior border is shifted anteriorly, and its posterior expression later extends throughout the entire opisthosoma. Besides these differences, the ring-like patterns in the developing walking legs differ tremendously between the two paralogs, with *Scr-1* showing expression in the L2-4 appendages, with stronger expression in L3, and more rings of expression per appendage. Scr-1 is first expressed in the center of the germ disc at stage 5. The position of the cumulus is marked by a ‘c’ (*A*). The posterior cap-like expression widens during stage 6. *Scr-1* now forms an open ring, localized roughly halfway between anterior rim of the opening germ disc and the future posterior center of the germ disc. The cells posterior to the ring also express *Scr-1*, but at a much lower level (*B*). At stage 7, the ring splits up into two stripes (asterisks) (*C*) and a bit later two new stripes appear (carets) (*D*). The most anterior stripe of expression lies between L1 and L2 at stage 8, the second anterior stripe lies between L2 and L3. The two posterior stripes cover L3 and L4. The weak expression of *Scr-1* continues posterior to these stripes (*E* and *F*). *Scr-1* is predominantly expressed in the limb buds and the ventral neuroectoderm of stage 9.1 embryos (*G*). The anterior expression border is directly posterior to the L1 limb bud. Expression of *Scr-1* continues in the opisthosoma, but is much weaker, except for a domain in the growth zone (arrowhead) (*G*). This expression in the growth zone continues to the end of segmentation (*H* and *I*). No more expression is visible at the posterior end of the embryo at stage 11 (white caret) (*J*). *Scr-1* expression forms multiple rings in the legs (*H-J*), but it is much more strongly expressed in L3 compared to L2 and L4. At stage 9 late the expression of *Scr-1* in the neuroectoderm can also be found in every segment of the opisthosoma (arrow in *H*). Each panel shows the same embryo viewed laterally (left) and ventrally (center and right). Anterior is to the left. Abbreviations: See Fig. S10.

**Figure S16. *P. tepidariorum Sex combs reduced-2* expression**

*Scr-2* expression appears later than *Scr-1* (Fig. S15), and it is restricted to a much smaller domain. A stripe of *Scr-2* expression first appears at stage 6 (*A*). The stripe broadens during stage 7 (*B*), at the end of which it starts to split into two stripes (carets), and a third new stripe appears posterior to the initial strips (asterisk) (*C*). At stage 8.1 there are four stripes that are located in the posterior parts of L2, L3, L4 and the newest stripe appears in O1 (white arrow) (*D* and *E*). In limb bud stages, only the L3 and L4 limb buds carry *Scr-2* expression, mostly in their distal tips. Expression in L2 is restricted to the neuroectoderm (not shown), and expression in O1 is restricted to the posterior part of each hemisegment as well (*F*). At later stages, *Scr-2* expression in L3 and L4 refines to several rings in the distal part of the legs. Expression in L2 becomes undetectable. (*G-I*). At stage 10.1, *Scr-2* is visible in a neuroectodermal patch in L4 (arrow) and a dot in O1 (arrowhead) (*H*). Each panel shows the same embryo viewed laterally (left) and ventrally (center and right). Anterior is to the left. Abbreviations: See Fig. S10.

**Figure S17. *P. tepidariorum fushi tarazu* expression**

*ftz* is first expressed at stage 6 in a semicircle at the posterior end where the SAZ is forming (arrow) (*A*). The bracket marks an anterior broad stripe of *ftz* expression. This domain of expression gets weaker during stages 7 and 8, where mostly only the anterior border of the domain anterior to L2 (arrowhead in *D*) is visible. The expression is almost invisible when it is viewed ventrally. The expression in the SAZ persists, and does not clear from the posterior end (*A-G*). It emanates stripes at the anterior end of the SAZ (arrow in *C-G*). After clearing from the posterior SAZ, stripes stay visible at the anterior border of the last formed segment (white arrow in *D* and *G*) until the next stripe of *ftz* expression forms. Only after segmentation finishes, the expression at the posterior disappears (arrow in *H*). Meanwhile, the anterior border of the Hox expression domain stretches from L2-L4 and is predominantly found ventrally (black arrowheads in *F* and *H*). The legs on one side of a stage 11 embryo were dissected off to show the ventral Hox domain, which has concentrated to one spot per hemisegment (black arrowheads) within the L2-L4 domain (*H*). At stage 10, *ftz* expression appears in a ring near the distal tip of L3 (carets in *G-H*). Each panel shows the same embryo, viewed laterally (left) and ventrally (right, or center and right in *E-H*). Anterior is to the left. Abbreviations: See Fig. S10.

**Figure S18. *P. tepidariorum Antennapedia-1* expression**

Expression of *Antp-1* appears earlier than *Antp-2* (Fig. S19) and it is initially only expressed in the presumptive O1 segment, but then broad expression is found in every segment added by the SAZ. *Antp-1* expression first develops in the SAZ at stage 7 (*A*). This expression transforms into a stripe and is followed by cells that show only weak expression of *Antp-1* (*B*), before new expression appears at the posterior end of the SAZ (*C*). The first two stripes can be allocated to the first two opisthosomal segments at stage 8 (*D*). New stripes keep appearing from the segment addition zone (arrows, *E-H*) until the end of segmentation (*G*). At stage 9, the anterior border of *Antp-1* expression reaches into the posterior half of L4 (arrowhead, *E*). Expression is now also seen in the ventral neuroectoderm, where it forms longitudinal rows in each hemisegment (white arrowheads, *E-G*), and rings in the L4 appendage (caret, *F* and *G*). Throughout development, *Antp-1* expression is always strongest in O1 and the anterior half of O2. Each panel shows the same embryo, viewed laterally (left) and ventrally (right, or center and right in *E-G*). Anterior is to the left. Abbreviations: See Fig. S10.

**Figure S19. *P. tepidariorum Antennapedia-2* expression**

In contrast to *Antp-1* (Fig. S18), *Antp-2* expression appears later, and is more restricted than *Antp-1*’s expression. While the strongest domain of *Antp-1* overlaps with *Antp-2*, the two paralogs differ in the exact pattern in the developing legs and the neuroectoderm and in difference to *Antp-1, Antp-2* later also expands into the prosoma. *Antp-2* first emerges in the first two opisthosomal segments at stage 8.2 (arrows) (*A*). The *Antp-2* domain expands anteriorly into the posterior part of L4 during stage 9 (white arrowheads) (*B-D*). Additionally, *Antp-2* forms rings of expression in the L4 appendage (carets in *C* and D). While initially the very posterior part of O2 is free of *Antp-2* expression (the O2/O3 border is demarcated by a black vertical line in *D-F*), during stage 10, expression can also be found at the O2/O3 border (arrows) (*F*). Furthermore during stage 10, the expression pattern refines, in that the L4 and O1 *Antp-2* expression is mostly in the ventral neuroectoderm and not found dorsally, while the O2 expression is excluded from the most ventral region (*E-G*). Starting at stage 10, *Antp-2* is also expressed in one dot each in the opisthosomal limb buds on O4 and O5 (arrowheads) (*E-G*). Each panel shows the same embryo, viewed laterally (left) and ventrally (right, or center and right in *E-G*). Anterior is to the left. Abbreviations: See Fig. S10.

**Figure S20. *P. tepidariorum Ultrabithorax-1* expression**

Expression of *Ubx-1* appears slightly earlier than *Ubx-2* (Fig. S21). The two *Ubx* paralogs share a similar expression domain, with the anterior border differing only by half a segment, however, the individual patterns within the segments expressing *Ubx* are unique and in some parts nonoverlapping. *Ubx-1* is first expressed at stage 8.1 in the SAZ and in a weak stripe anterior to that (arrow) (*A*). This stripe gets stronger and the posterior domain enlarges (*B* and *C*) so that all new tissue added posteriorly expresses *Ubx-1* at equal levels (*D-F*). The anterior border is located in O2, while the very anterior part of O2 initially does not express *Ubx-1* (segmental boundaries in *C-F* indicated by black vertical lines) until the end of stage 9.2, when it is expressed dorsally and in the neuroectoderm also in the anterior part of O2 (white arrow in *E*). Each panel shows the same embryo, viewed laterally (left) and ventrally (right). Anterior is to the left. Abbreviations: See Fig. S10.

**Figure S21. *P. tepidariorum Ultrabithorax-2* expression**

*Ubx-2* is first expressed slightly later than *Ubx-1* (see Fig. S20) at stage 8.2, in the SAZ, as well as in two weaker stripes in the O3 and O4 segments (arrows) (A). Expression is initially restricted to the posterior part of O3 (B), but eventually the anterior expression border extends ventrally into the posterior part of O2 (*C-F)*, and is also found in the posterior part of the O2 limb bud (caret in *E* and *F*). Expression is strongest in O3, but otherwise is fairly uniform in all segments posterior to this one (*D-F*). Each panel shows the same embryo, viewed laterally (left) and ventrally (right). Anterior is to the left. Abbreviations: See Fig. S10.

**Figure S22. *P. tepidariorum abdominalA-1* expression**

*abdA-1* expression appears only marginally earlier than *abdA-2* (Fig. S23), and *abdA-1’s* anterior expression border is more dynamic than the one of *abdA-2*, and eventually extends into the posterior half of O2. *abdA-1* expression starts in the SAZ during stage 9.1, posterior to O5 (A and B). By stage 10.1, the anterior border of *abdA-1* expression has extended into the posterior part of O3 (arrowhead, *C*). While the dorsal border of *abdA-1* expression remains there, in the ventral neuroectoderm the expression expands anteriorly into the posterior half of O2 (arrowhead, *D*). These borders of expression persist during later development (*E* to *G*). Each panel shows the same embryo, viewed laterally (left) and ventrally (right). Anterior is to the left. Abbreviations: See Fig. S10.

**Figure S23. *P. tepidariorum abdominalA-2* expression**

*abdA-2* expression emerges only very slightly later than *abdA-1* (Fig. S22), however, in contrast to *abdA-1, abdA-2*’*;s* anterior expression domain remains more static and stays within O4 throughout development. *abdA-2* expression first appears at stage 9.1 in the SAZ (*A*). Slightly later, shortly before the opisthosomal limb buds appear, *abdA-2* additionally emerges in the posterior part of O4 (*B*). From stage 9.2 onwards, *abdA-2* is expressed in the entire O4 segment and all segments posterior with strong expression in the opisthosomal limb buds, (*C-F*). Each panel shows the same embryo, viewed laterally (left) and ventrally (right). Anterior is to the left. Abbreviations See Fig. S10.

**Figure S24. *P. tepidariorum AbdominalB-1* expression**

*AbdB-1* is first expressed much later than *AbdB-2* (Fig. S25) in O6 and in a stripe in the anterior-most portion of the segment addition zone (*A*). The anterior expression border subsequently shifts anteriorly, first into the posterior part of O5 (arrowhead) (*B* and *C*), and the dorsal part of the *AbdB-1* domain then shifts into the posterior portion of O4 (arrowhead) (*D* and *E*). From the beginning of opisthosomal limb bud development, it is strongly expressed in the O5 limb buds. At stage 9.2, the ventral part of the *AbdB-1* domain expands into the posterior half of O2 (arrow in *D*). These anterior borders remain the same throughout the rest of embryonic development (*E-G*). Each panel shows the same embryo, viewed laterally (left) and ventrally (right, or center and right in *D-G*). Anterior is to the left. Abbreviations: See Fig. S10.

**Figure S25. *P. tepidariorum AbdominalB-2* expression**

In contrast to *AbdB-1* (Fig. S24), *AbdB-2* shows a very dynamic expression pattern. Moreover, if one takes Hox gene collinearity rules into account, whereby genes located more posteriorly in the Hox gene cluster are expressed later during development, *AbdB-2* does not follow this rule. Instead, *AbdB-2* expression emerges with the appearance of the segment addition zone at stage 6 (*A*). It is continuously expressed in the segment addition zone until the end of segmentation (*B-H*). Additionally, from stage 9.1 on, weak *AbdB-2* expression is found posterior to the O3/O4 border (arrowhead) (*F*). Slightly later, additional expression appears in the O2 limb buds, in the prospective genital opening (arrow) (*G* and *H*). Dorsally, the *AbdB-2* expression domain extends from O5 to the posterior end (*H-J*), but in the ventral neuroectoderm, the *AbdB-2* expression border is located in the posterior half of O3 (arrowhead in *I*). The vertical line delineates the O2/O3 border, which the *AbB-2* expression does not reach in the neuroectoderm (*J*). Each panel shows the same embryo, viewed laterally (left) and ventrally (right, or center and right in *G-J*). Anterior is to the left. Abbreviations: See Fig. S10.

